# Genome Sequencing of Chinese yam (*Dioscorea polystachya*): analysis of PEBP gene family diversity and identification of a potential tuber inducing factor

**DOI:** 10.1101/2025.03.10.641988

**Authors:** Tatjana Ried, Marie Bolger, Jenny Riekötter, Toshiyuki Sakai, Janina Epping

**Affiliations:** Cologne, Germany; Institute of Bio- and Geosciences (IBG-4: Bioinformatics), Bioeconomy Science Center (BioSC), CEPLAS, Forschungszentrum Jülich GmbH, 52425 Jülich, Germany; Institute of Plant Biology and Biotechnology, University of Muenster, 48143 Muenster, Germany; Crop Evolution Laboratory, Kyoto University, Mozume, Muko, Kyoto 617-0001, Japan

**Keywords:** Chinese yam, whole genome sequencing, FLOWERING LOCUS T, tuber formation, *Solanum tuberosum*

## Abstract

Phosphatidylethanolamine-binding proteins (PEBPs) are a class of ancient plant proteins involved in flowering, seed development and storage organ formation. The FLOWERING LOCUS T subgroup was shown to be involved in the initiation of flowering and tuberization. We aimed to characterize the PEBP family in the tuber crop Chinese yam (*Dioscorea polystachya*) and to identify FLOWERING LOCUS T (FT) homologs potentially involved in Chinese yam tuber formation. Based on a *de novo* genome assembly of Chinese yam, we investigated members of the PEBP family *in silico*. Expression analysis of FT homologs in yam tuber tissue revealed DpFT3a as a candidate for a tuber inducing FT. We then performed functional characterization of DpFTs by localization and interaction studies, as well as constitutive overexpression in *Arabidopsis thaliana* and *Solanum tuberosum*. Analysis of the genome annotations revealed members of the PEBP-family in Chinese yams. Identified PEBPs were subjected to phylogenetic analysis and transcriptomic studies. Of the identified 9 FT homologs, DpFT3a, which was strongly expressed in yam tubers, was found to cause a severe tuberizing phenotype in transgenic potato (*Solanum tuberosum* cv. Désirée), indicating a role of DpFT3a in storage organ formation. We identified a Chinese yam specific FT homolog and provide strong indication for a role of this FT in tuber formation in yam as well as its potential to induce tuberous structures in potato. Our results complement the current knowledge on PEBP proteins in plants.

## Introduction

Geophytes are plants that are able to produce storage organs in the form of tubers, corms, bulbs or rhizomes for reproduction and endurance of adverse environmental conditions (Khosa et al. 2021). A number of these storage organs serve as an important food source for calories and nutrients; they are produced by plant species such as the potato, sweet potato, carrot, cassava and yam. To date, most research concerning storage organ formation has been done in potato (*Solanum tuberosum*), the most important tuber crop globally (Xu et al. 2011). Several molecular factors were found to participate in tuberization, and special attention was paid to SELF PRUNING 6A (StSP6A), a FT homolog, which plays a crucial role in tuber induction in potato (Navarro et al. 2011). Initially, FT was shown to play a critical role in regulating floral induction in *Arabidopsis thaliana* as well as many other angiosperms (Turck et al. 2008; Pin et al. 2010). Dependent on day length, the transcription factor CONSTANS (CO) activates *FT* expression in the phloem companion cells in *Arabidopsis* leaves (An et al. 2004; Wigge et al. 2005). Several studies showed that protein and mRNA of FT are phloem-mobile and traffic through the phloem to the shoot apical meristem (SAM) (Corbesier et al. 2007; Lin et al. 2007; Tamaki et al. 2007; Yoo et al. 2013). In the SAM, FT interacts with the transcription factor FLOWERING LOCUS D (FD), mediated by 14-3-3 proteins, to induce the expression of floral identity genes, turning the SAM into an inflorescence meristem (IM) (Wigge et al. 2005). TERMINAL FLOWER 1 (TFL1), a protein with a high amino acid similarity to FT, acts antagonistically to FT by interacting with FD to repress floral identity genes at the tip of the IM. Thus, TFL1 maintains the meristemal identity of the IM, while floral meristems form at the sides (Koornneef et al. 1991; Shannon and Meeks-Wagner 1991; Shannon and Meeks-Wagner 1993; An et al. 2004; Abe et al. 2005; Lee et al. 2008; Taoka et al. 2011). In tobacco (*Nicotiana tabacum*) and sugar beet (*Beta vulgaris*), antagonistic FTs in floral development were identified (Pin et al. 2010; Harig et al. 2012). FTs with variations in the amino acid sequence of functionally conserved motifs, act as flower-repressing factors besides TFL (Pin et al. 2010; Harig et al. 2012). Both FT and TFL are members of the PEBP protein family, defining two of three subclades in angiosperms (FT-like, TFL-like and MFT-like) (Kardailsky et al. 1999; Kobayashi 1999).

Tuber induction in the strictly short day (SD) tuberizing potato subspecies *Andigena* was found to be similar to the flower inducing process of other plants (Navarro et al. 2011). Similar to flower development, a CO homolog in potato (*StCO*) is expressed higher under long days (LDs), but instead of inducing the expression of the tuber-inducing *StSP6A*, it induces the expression of the FT homolog *StSP5G*. In the leaf, StSP5G prevents tuberization by inhibiting *StSP6A* expression. As StCO is unstable under SD conditions, *StSP5G* expression is not induced and *StSP6A* is not repressed. The StSP6A protein moves through the phloem to the stolon, a modified underground shoot that develops into a tuber (Martinez-Garcia et al. 2002; González-Schain et al. 2012; Abelenda et al. 2016; Sharma et al. 2016). In the stolon, StSP6A interacts with FD, mediated by 14-3-3 proteins. This interaction results in the formation of the so-called tuber-activation complex (TAC), regulating the expression of tuber formation genes (Teo et al. 2017). FT involvement in storage organ formation was also investigated in the monocotyledonous crop onion (*Allium cepa*). Here, AcFT homologs were shown to initiate or inhibit bulb induction and regulate flowering time, dependent on daylength and vernalization (Lee et al. 2013).

Tuber crops play an important role in human nutrition and global food security (Scott et al. 2000). A profound understanding of tuber development in these crops is a prerequisite for improved breeding strategies. One such important tuber crop is yam (*Dioscorea spp*.). Species commonly known as yams belong to the monocotyledonous genus *Dioscorea*. Yams are mostly annual, dioecious plants that produce starchy underground tubers, which emerge from the hypocotyl (Coursey 1967; Shewry 2003). Although mostly unknown in Europe, they are cultivated as staple crops, predominantly in Western Africa, but also Asia, Southern and Central America and the Pacific Islands (Irvine 1952; Ayensu and Coursey 1972; Opara 2003; Bhattacharjee et al. 2011; Tamiru et al. 2017).

Chinese yam (*Dioscorea polystachya*) is cultivated in temperate climates, primarily Japan and China (O’Sullivan J. N. 2010). The tubers of Chinese yam show a geotropic response in the distal end. While the distal end grows downwards and thickens, the proximal end stops growing and hardens (Shewry 2003). Besides serving as a carbohydrate source, Chinese yam has many beneficial health ingredients. In several studies, whole tubers or separate compounds were shown to, amongst others, increase insulin sensitivity, suppress fat uptake and exhibit cholesterol-lowering effects (KWON et al. 2003; Gao et al. 2007; Liu et al. 2009; Son et al. 2014; Fan et al. 2015). Despite its importance as a food crop and health benefits, yam is considered an orphan crop (Otoo 2017; Tamiru et al. 2017; Mabhaudhi et al. 2019). Challenging conditions such as labor-intensive cultivation, a long growth cycle, sparse and asynchronous flowering as well as prevalent asexual propagation slow down breeding attempts (Burkill 1960; Hahn, S. K. Osiru, D. S. O. Akoroda, M. O. and Otoo 1987; Asiedu et al. 1997; Mignouna et al. 2003; Mignouna et al. 2007). Furthermore, yams are polyploid and highly heterozygous, impeding investigations of the genetic background of tuber growth and traits (Mignouna et al. 2003).

To improve our understanding of tuber development in yam, we investigated the PEBP family in Chinese yam with the aim to identify FTs potentially involved in the tuberization process. To identify PEBP homologs *in silico*, we generated and assembled a *de novo* whole genome assembly using PacBio Hifi reads. Intra-species ploidy variation has previously been described in many species of *Dioscorea* with diploid, triploid and tetraploid levels reported for *D. alata* (Arnau et al. 2009), a species closely related to Chinese yam. Two different cultivars of Chinese yam have been reported as having a chromosome number of 100 and 140 (Babil et al. 2013). Given that the basic chromosome number for *D. alata* is reported as 20 (x=20), we expect Chinese yam to be pentaploid or heptaploid. In the present study, we used the *de novo* genome data to characterize the PEBP family *in silico*. Based on sequence analyses, we selected a FT homolog potentially involved in tuber formation and show its functionality as a potential tuber inducing protein *in planta*.

## Materials and Methods

### *De novo* Genome sequencing

Tubers of *D. polystachya* were provided by a local farmer (St. Calude de Diray, Loir-et-Cher, France) from a variant we titled ‘F60’ (Riekötter et al. 2023). F60 plants grown from seed tubers were cultivated in raised bed (1.2 × 0.8 × 1.4 m) in Münster, Germany (51°57’55.3”N 7°36’54.1”E) between April and September 2020. Leaf material of young leaves was flash-frozen in liquid nitrogen and sent to DNA Sequencing Center at Brigham Young University, Utah, US for DNA extraction and PacBio SMRT sequencing. Sequencing was performed on a PacBio Sequel II using three 8M SMRT cells and 30 hour collection time.

### Genome assembly

Raw PacBio circular consensus sequencing (ccs) reads were converted to HiFi fastq files using the PacBio pipeline. The assembly of the genome was performed using the Hifiasm assembler version 0.17.6. (Cheng et al. 2021) specifically designed to handle long and accurate PacBio HiFi reads. Hifiasm employs an overlap-layout-consensus (OLC) strategy to construct a *de novo* assembly. Structural annotation of the genes was performed using Helixer, a novel state-of-the-art ab-initio gene caller which uses deep neural networks to structurally annotate genes in a genome (Holst et al. 2023). The resulting output was assessed using BUSCO and compared to the results of BUSCO on the genome.

### *In silico* analysis of DpPEBP sequences

Protein function annotation was assigned to protein sequences using the online tool Mercator4 v5.0 (Schwacke et al. 2019; Bolger et al. 2021) (Table S1). Multiple sequence alignment (MSA) of PEBP protein sequences was performed using the MUSCLE algorithm. A phylogenetic tree was constructed by a Neighbor-Joining method based on *p*-distance with 1000 bootstrap replicates using the MEGA11 software (Tamura et al. 2021). PEBPs from species other than *D. polystachya* had the following NCBI GenBank accession numbers: *Arabidopsis thaliana* (At), BAA77838.1, AAD37380.1, BAB10165.1, AAB41624.1, AAF03937.1; *Beta vulgaris* (Bv), ADM92608.1, ADM92610.1; *Nicotiana tabacum* (Nt), JX679067, JX679070, AAD43528.1; *Oryza sativa* (Os), BAB61028.1, BAB39886.1, *Solanum tuberosum* (St), BAV67096.1, XP_006365457.1, NP_001274897.1, *Triticum aestivum* (Ta), BAK78895.1. MSA of FT protein sequences was visualized using the Jalview v2.11.2.7 program (Waterhouse et al. 2009). Visualization of gene structure of *DpPEBP* genes was performed with TBtools v1.123 (Chen et al. 2020).

### Expression of PEBPs in tuber tissue

Expression of *PEBP* family members in Chinese yam tubers were analyzed using the recently published transcriptomic data of *D. polystachya* var. F60 (Riekötter et al. 2023) (Accession number: PRJNA942579). Raw sequencing data were filtered using fastp v0.12.4 to cut adapter sequences and to remove bad quality reads (Chen et al. 2018). Mapping of clean reads against the genome of *D. polystachya* var. F60 was performed using HISAT2 v 2.2.1 with following parameters: -k 1 --no-mixed --no-discordant (Liao et al. 2014; Kim et al. 2015). Counting of mapped reads per gene was performed using featureCounts v2.0.1 (Liao et al. 2014).

### Generation and analysis of transgenic *S. tuberosum* plants

The coding sequence of DpFT3a was amplified from cDNA of F60 tuber tissue using primers DpFT3_NcoI_fwd and DpFT3_XbaI_rev (Table S2) deduced from the genomic sequence. Amplified fragments were purified using agarose gel electrophoresis and the Gel and PCR clean-up-Kit (Macherey-Nagel, Düren, Germany) and ligated into plant expression vector plab12.10-Q35S (Xing et al. 2014; Stolze et al. 2017) for overexpression in *S. tuberosum* cv. Désirée resulting in plab12.10-Q35S-DpFT3a. The plab12.10-Q35S-DpFT3a was stably transformed into *S. tuberosum* cv. Désirée as described previously (Hoekema et al. 1983; Horsch et al. 1985) using the *Agrobacterium tumefaciens* strain LBA4404. Transformed leaves were cultivated in sterile culture on selective solid medium (4.4 g L^-1^ Murashige and Skoog salt solution (MS) including vitamins, 20 g L^-1^ sucrose, and 6.5 g L^-1^ agar, 75 mg L^-1^ kanamycin, pH 5.8), in sterile environment under long day conditions (16 h artificial light (100 μmol m^-2^s^-1^)/ 8 h darkness) at 24/17 °C. Regenerated plants were tested for transgene integration. Regenerated transgenic plants were moved from tissue culture to the greenhouse upon rooting. For expression studies of transgenes, plant material was frozen in liquid nitrogen immediately after harvest and RNA was extracted using innuPrep Plant RNA Kit (IST Innuscreen, Berlin, Germany) with DNaseI-digest as described by the manufacturer. Subsequently, 500 ng of isolated RNA were transcribed into cDNA using the PrimeScript RT Master Mix (Takara Bio Europe, Saint-Germain-en-Laye, France) accordingly to the manufacturer’s instructions. For quantitative real-time PCR the SYBR™ Green PCR Master Mix (KAPA Biosystems, Wilmington, Massachusetts, USA) and the primers DpFT3_qPCR1_fwd/ DpFT3_qPCR1_rev and qRT_StGAPDH_fw/ qRT_StGAPDH_rev (Table S2) were used. A 10 μL reaction mixture containing 5 μL KAPA SYBR FAST qPCR Master Mix (2×) Kit (Roche, Basel, Switzerland), 2.5 μL cDNA (diluted 1:5 or 1:10) and 2.5 μL primer mix (2 μM) was prepared and qRT-PCR was performed on a CFX96 Touch Real-Time PCR Detection System (Bio-Rad Laboratories, Hercules, CA, USA) with following program configuration: 95°C for 3 min, followed by 44 cycles of 95°C for 3 s, 60°C for 20 s and 95°C for 5 s. Melt curve analysis was performed in 0.5°C increments from 58 to 95°C to confirm gene specific amplification. Reactions were carried out in technical triplicates for each biological replicate. The housekeeping gene *StGAPDH* served as internal control and was used for gene expression normalization. Relative gene expression was calculated using the 2^-ΔΔCt^ method (Livak and Schmittgen 2001).

Further methods can be found in the Supplementary material.

## Results

### *De novo* genome sequence

The quality of the assembled genome was assessed using several metrics, including N50, assembly length, and number of contigs (Table 1). Due to the high ploidy of the *D. polystachya* genome, hifiasm struggled to create a high quality, contiguous genome despite the use of high quality PacBio Hifi reads. The draft assembly was however assessed as satisfactory for the purpose of gene exploration. The completeness of the genic component was assessed with BUSCO (Simão et al. 2015; Manni et al. 2021) and showed a 98% overall completeness when using the embryophyta_odb10 dataset.

**Table 1.** Quality of *de novo D. polystachya* genome assembly.

|  |  |
| --- | --- |
| Assembly name | yam_v0.16.1.bp.p_ctg |
| Length (bp) | 1,807,892,738 |
| Number of contigs | 2,462 |
| Contig N50 length (bb) | 9,966,655 |
| BUSCO (embryophyta_odb10) | C:98.7%[S:1.3%,D:97.4%],F:0.6%,M:0.7%,n:1614 |

### The PEBP-family has 75 members in Chinese yam

In plants, the PEBP-family is divided into the three subclades of MFT-like, TFL-like and FT-like, based on their function and conservation of essential motifs (Kardailsky et al. 1999; Kobayashi 1999). To gather information about the PEBP-family in Chinese yam, the functional annotation of the genome was screened for members of this family. Overall, 75 genes with annotation as PEBP were found based on the functional annotation obtained by Mercator4 (with prot-scriber and swissprot enabled) and their similarity to reference sequences from known PEBPs (Table S1). Based on the annotations and phylogenetic analysis (Fig. 1), the DpPEBPs were grouped into 22 MFT-like genes, 19 TFL-like genes and 34 FT-like genes and named and grouped accordingly. The examination of the predicted intron-exon structure of *DpPEBP* genes revealed the presence of four introns in most of the *PEBP* genes (Fig. S1).

**Fig. 1.**
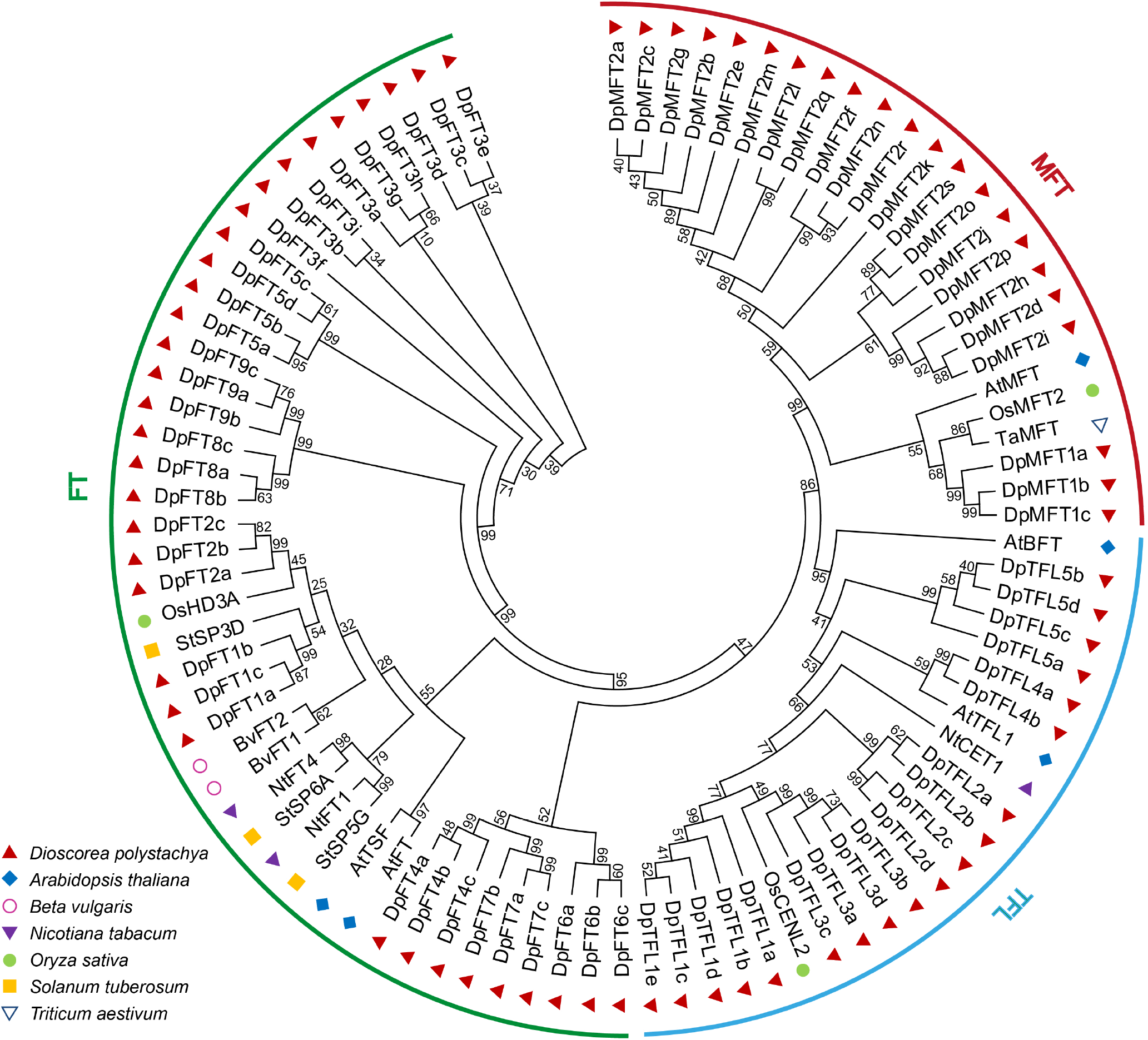
The PEBP family in *D. polystachya*. Phylogenetic analysis of all 75 identified *DpPEBP* genes in the *D. polystachya* genome plus sequences from PEBPs with known functions from other plant species. Protein sequences were aligned using MUSCLE algorithm. Phylogenetic tree was generated using Neighbor-Joining method with 1000 bootstrap replication. Bootstrap values are shown at each node in percent.

A phylogram of DpPEBPs and known PEBP protein sequences of several other mono- and dicots was created to investigate similarities of DpPEBPs with PEBPs with known functions (Fig. 1). In the MFT-like subclade, DpMFT1 is close to the MFTs of the rice, wheat and Arabidopsis, while DpMFT2 showed greater distance. DpTFL1, DpTFL2 and DpTFL3 grouped very close with OsCENL2, a TFL-like homolog, while DpTFL4 was closest to Arabidopsis AtTFL1 and DpTFL5. Within the FT-like subclade, only DpFT1 and DpFT2 showed high similarity to known FT homologs. All other DpFTs are grouped further apart indicating lower sequence similarity to plant FT homologs described in literature.

### Five *DpFTs* are expressed in tuber tissues

Several studies have revealed the importance and function of amino acid motifs of FT-proteins (Hanzawa et al. 2005; Ahn et al. 2006; Ho and Weigel 2014). We generated an amino acid alignment of DpFTs and FT homologs of known activating or repressing function (Fig. 2). The best studied motifs of FTs are REY, YAPGWRQ and LYN (highlighted in Fig. 2). The Y residues of the REY and LYN motif are conserved in all DpFTs, except for DpFT5, which showed the characteristic REH motif for TFLs. The YAPGWRQ motif, described in inducing FTs, was present in DpFT1-3 and DpFT5. In DpFT4, DpFT6, DpFT7, DpFT8 and DpFT9, alterations of the YAPGWRQ were detected, similar to FTs of repressing function (Pin et al. 2010; Harig et al. 2012). Taken together, DpFT1, DpFT2, DpFT3 and DpFT5 contained all of the amino acid motifs, described as essential for a flower- or tuber-inducing function. Next, we investigated the gene expression of potential tuber-inducing DpFTs in Chinese yam tuber tissue.

**Fig. 2.**
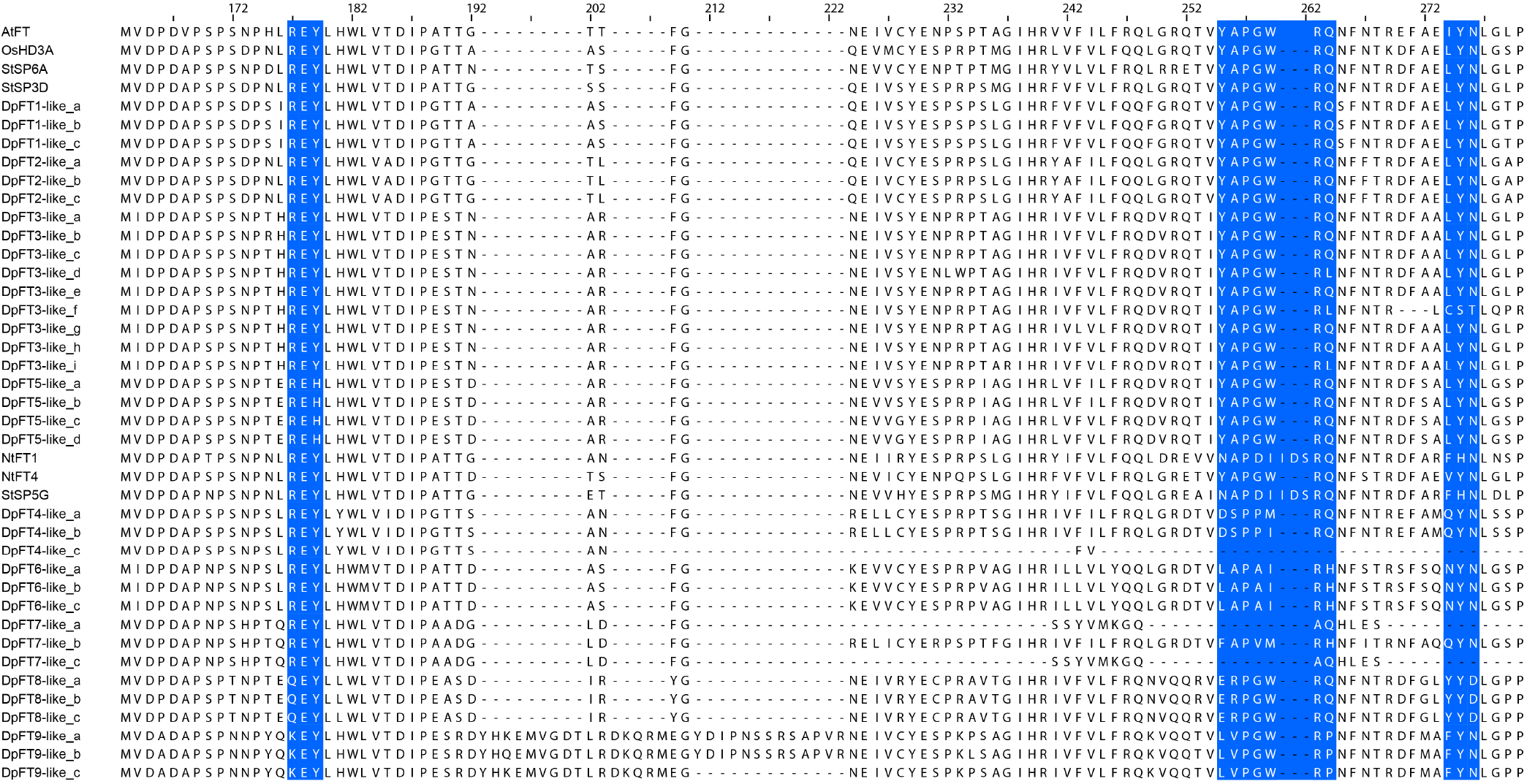
Partial amino acid alignment of FT homologs. DpFTs were aligned to inducing and repressing FTs from Arabidopsis, rice, tobacco and potato. Conserved motifs are highlighted in blue. The alignment was performed using MUSCLE algorithm.

Based on previously published Chinese yam transcriptome data (Riekötter et al. 2023), we calculated the normalized read counts of all *DpPEBPs* (Liao et al. 2014). Expression was analyzed in the tuber head, middle and tip, to reveal differences in the expression pattern along the tuber growth gradient, as the tuber tip is the actively growing tuber part, while the tuber head is the oldest tuber part (Fig. 3). Differences in expression profiles were observed between the identified *DpFTs*. Highest expression was detected for variants of *DpFT7*, followed by *DpFT3* and *DpFT5* variants, respectively. All the three *DpFTs* expressed in tuber tissue showed highest expression in the head, declining towards the tip. Of *DpMFTs* only *DpMFT1* was expressed in tuber tissue with *DpMFT1b* showing the highest expression in the tuber head. All identified *DpTFLs* were expressed in tuber tissue with *DpTFL4b* showing high expression in the tuber head, declining toward the tip.

**Fig. 3.**
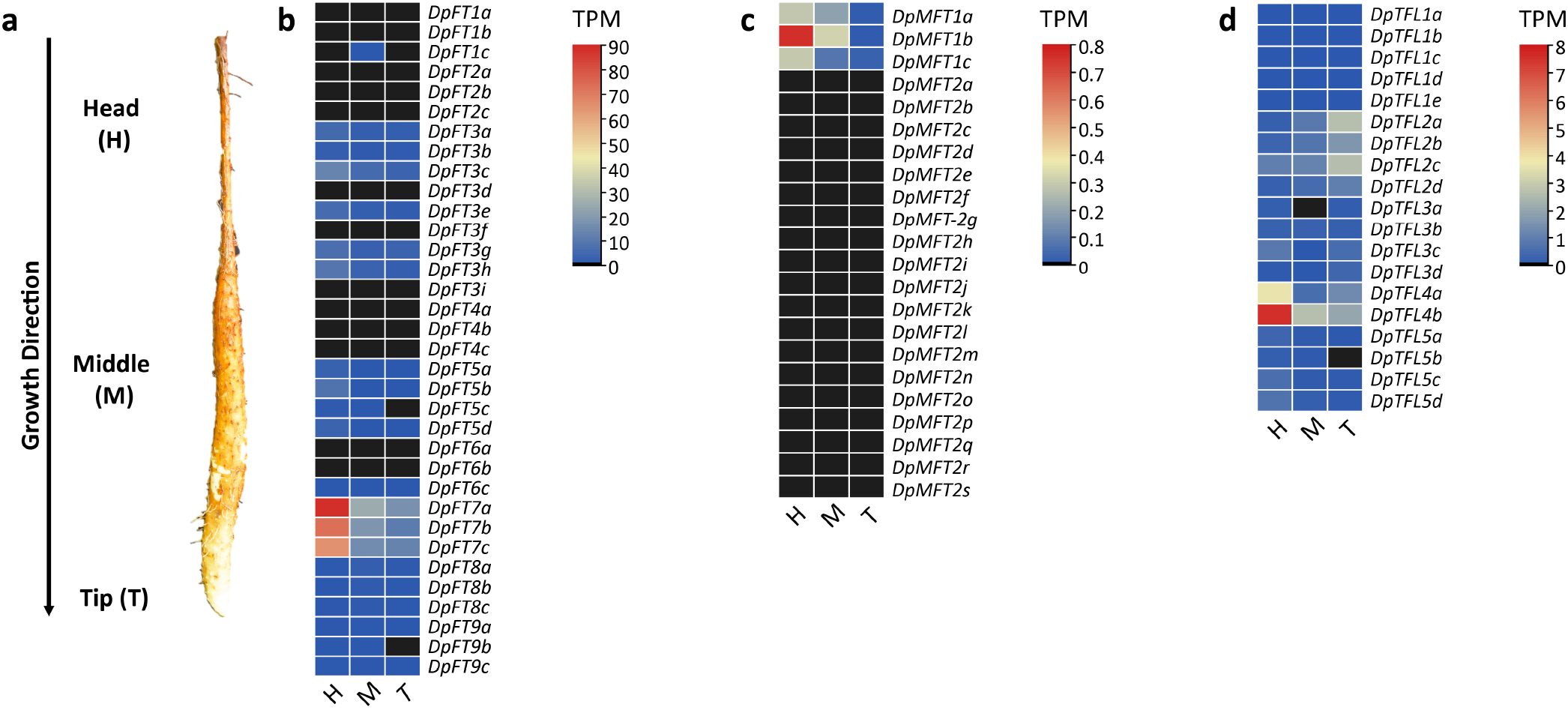
Expression of DpPEBPs in tuber tissue. Expression levels were calculated in the head (H), middle (M) and tip (T) region of a growing tuber as shown in (a). Based on tuber transcriptome data (Riekötter et al. 2023) the expression levels of DpFTs (b), DpMFTs (c) and DpTFLs (d) were analyzed. TPM: transcripts per million (Raw data can be found in Table S3).

### DpFT3a is a functional FT with inducing function

In search of a tuber inducing yam FT, we then focused on DpFT3a, since it contains all conserved motifs of inducing FTs and was expressed in tuber tissue. Interaction of FT with FD and the generation of a TAC is reported to be vital for the inducing and repressing functions of FT-like proteins, which also implies that the proteins must not be secluded within different compartments of the cell. Hence, we studied the interaction of DpFT3a with a potential DpFD protein, that we named DpFD1, which was isolated from tuber tissue (Table S1). Cellular localization of both proteins was also performed. Both approaches were investigated upon transient expression in *N. benthamiana* leaves. To monitor DpFT3a and DpFD1 localization within cells, the N-terminal region of DpFT3a and DpFD1 were fused with the fluorophore Venus. A previous study showed that FD localizes to the nucleus, while FT localizes to the nucleus and the cytoplasm. Upon protein interaction, the FD-FT complex localizes to the nucleus (Abe et al. 2005). DpFD1 was indeed found to localize to the nucleus, while DpFT3a localized to the nucleus and the cytoplasm in *N. benthamiana* epidermal cells (Fig. S2).

Next, we investigated the capability of DpFT3a to form a protein complex with DpFD1. Here we performed a BiFC assay, in which the N-terminal part of mRFP was fused to the N-terminus of DpFT3a and the C-terminal part of mRFP was fused to the N-terminus of DpFD1, resulting in red fluorescence upon interaction of DpFT3a and DpFD1. Detection of fluorescence of the reporter protein in the nucleus indicated interaction between DpFD1 and DpFT3a (Fig. S3). Furthermore, we investigated the function of DpFT3a upon overexpression in *A. thaliana*. Systemic overexpression of *DpFT3a* in *A. thaliana* plants resulted in significantly earlier flowering of transgenic lines compared to wild type (WT) control plants, indicating that DpFT3 is a functional FT with an inducing function *in planta* (Fig. S4).

### DpFT3 induces a tuberous phenotype in potato

To investigate whether DpFT3a influences storage organ formation, *DpFT3a* was overexpressed in *S. tuberosum* cv. Désirée under control of the CaMV 35S promoter. Several transgenic lines started to form tuberous organs during regeneration in tissue culture and showed strongly impaired shoot, leaf and root growth (Fig. 4 a-c). The produced tuberous structures had an elongated shape and Lugol-staining revealed starch accumulation in these organs (Fig. 6d-g). Consequently, these plants could not be transferred to the greenhouse and stayed in tissue culture (TC-lines). After removal from tissue culture, the tuberous organs did not sprout. Other lines could be transferred to the greenhouse (GH-lines) and showed no phenotypic differences compared to WT plants (Fig. S5, Table S2). Expression analysis on leaf tissue in tissue culture showed that TC lines had an overall significantly higher *DpFT3a* expression, indicating that the phenotype was dosage dependent (Fig. 4h).

**Fig. 4.**
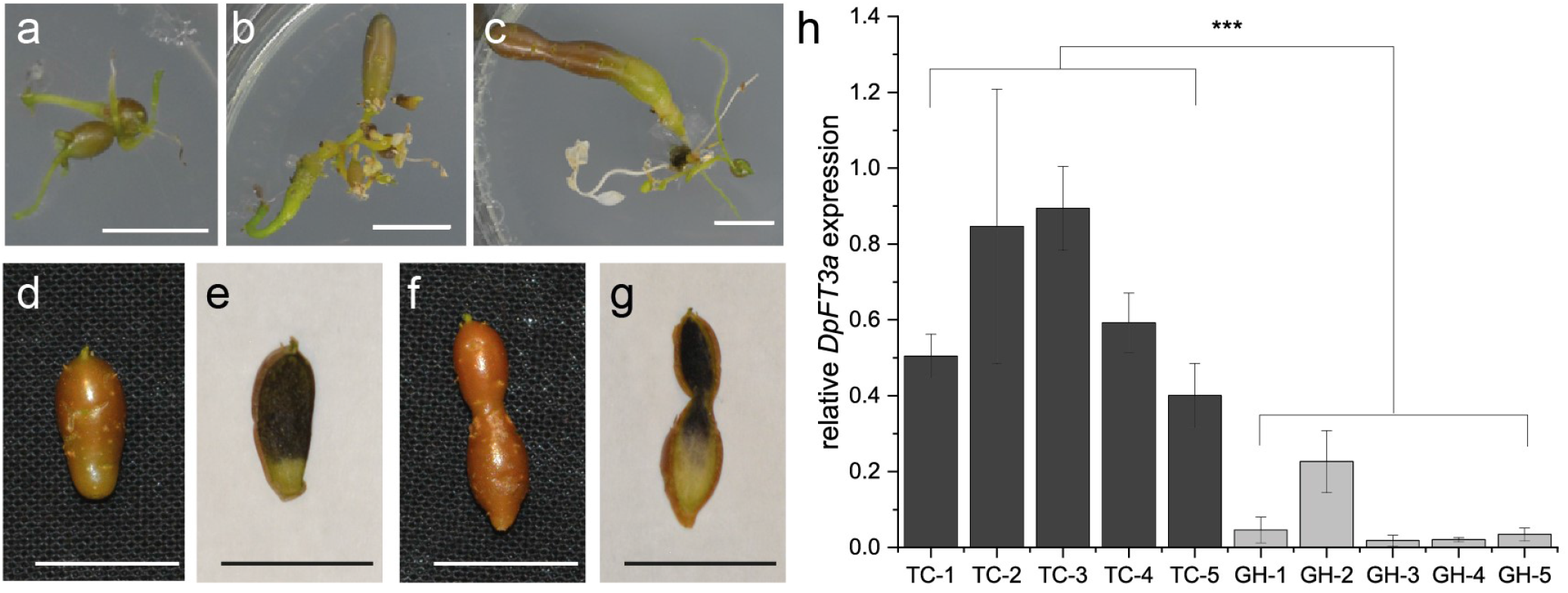
Overexpression of *DpFT3a* induces tuberous organs in potato. Transgenic potato lines with *DpFT3a* overexpression showed a tuberous phenotype with tuberous structures instead of shoots and produced neither functional roots nor leaves. Exemplarily, this phenotype is shown for three independent lines, TC-1 (a), TC-2 (b) and TC-4 (c) in tissue culture. Lugol-staining of the tuberous structures showed the accumulation of starch (d-g). d and e show the tuberous organs before cutting and e and g show the organs cut in half and stained with Lugol-staining. Scale bar = 1 cm. The observed phenotypes appeared in a dosage-dependent manner (h). While lines with high *DpFT3a* expression (TC) showed the tuberous phenotype, lines with lower *DpFT3a* expression levels (GH) could be transferred to the greenhouse and showed no phenotypic differences compared to WT plants. Shown is the relative *DpFT3a* expression in leaf tissue of five independent TC and GH lines regenerated in tissue culture (n=5). Difference in gene expression was calculated between all TC-lines and GH-lines via Wilcoxon ranked sum test. Asterisks indicate statistical significance: *** p-value ≤ 0.001.

## Discussion

### Genome sequence of *D. polystachya*

Despite the use of high-quality long reads, assembling of the genome to chromosome scale was not possible with the current data and available assemblers. We can see that the *D. polystachya* has a high overall BUSCO score which is comparable to *D. alata* (Bredeson et al. 2022) but with an extremely high level of duplication. On closer examination of the BUSCO duplication data, we can see that the gene duplication is largely 4x (which would roughly correspond to the 4x genome size versus *D. alata*). Since the main hifiasm assembly typically merges parental chromosomal copies, this suggests that the genome is likely octoploid (2n=8x=160). Upon checking the hifiasm haplotype phased assemblies, which try to keep parental copies separate, a roughly 7x in both BUSCO copy number and overall genome size was observed, indicating that cultivar F60 is septaploid (2n=7x=140) as has previously been reported for this species (Babil et al. 2013).

### Diversity of PEBP-family in Chinese yam

In the present study we investigated the genetic diversity of PEBPs in Chinese yam on the basis of a *de novo* genome sequence. Overall, 75 PEBP sequences were found in the genome. Sequence alignment and phylogenetic studies enabled grouping of PEBP genes into 22 MFT-like, 19 TFL-like and 34 FT-like genes. The classification of PEBPs into the subclades (MFT; TFL and FT) is in concurrence with previous studies on PEBP families in plants (Kardailsky et al. 1999; Chardon and Damerval 2005; Karlgren et al. 2011). The high diversity of PEBPs and the relative number of PEBPs in each group found in *D. polystachya* fits in with species analyzed in other studies (Chardon and Damerval 2005; Zheng et al. 2016; Dong et al. 2020; Zhang et al. 2021).

Furthermore, the relative distribution of the subgroups found in our study is in accordance with studies in other monocotyledonous species, which showed that monocots have a comparable number of MFTs and TFLs, but a higher number of FTs compared to dicots (Chardon and Damerval 2005; Karlgren et al. 2011; Zheng et al. 2016). This is also supported by recent finding in the White yam *D. rotundata*, where 10 PEBPs (8 FT, 1 TFL, 1 MFT) were identified in the genome (Susila and Purwestri 2023). The high number of FTs compared to TFL and MFT is probably caused by positive selection and diversification of FT function (Zheng et al. 2016). Phylogenetic analyses showed that FTs have diversified into up to 12 individual clades with 5 main lineages. *Dioscoreales* were shown to have three of these groups (Bennett and Dixon 2021), fitting with our phylogenetic analysis on DpFTs, which showed that DpFTs separate into three different groups. The study further suggests that the high number of FTs is thought to be the result of neofunctionalization of FT in developmental processes (Bennett and Dixon 2021).

### Phylogenetic relationship to known PEBPs

We analyzed the phylogenetic relationship of DpPEBPs to functionally characterized PEBPs from other plant species. DpFT1 and DpFT2 grouped with StSP3D and OsHd3a, both known to induce flowering in potato and rice respectively (Tamaki et al. 2007; Navarro et al. 2011), but not with other Solanaceae FTs. This is consistent with Chardon and Damerval, who hypothesized that PEBPs have undergone independent evolution within species (Chardon and Damerval 2005). In search for a potentially tuber-inducing FT (“tuberigen”), we took closer look at the protein motifs described to be relevant for their function in known plants. Four DpFTs (DpFT1, DpFT2, DpFT3, DpFT5) contained the conserved YAPGWRQ motif, which was found in several studies to be crucial for a flower- or tuber-inducing function (Ahn et al. 2006; Pin et al. 2010; Navarro et al. 2011; Harig et al. 2012). In onion, it was found that the bulb promoting AcFT1 has the slightly altered YAPnWRQ motif (Lee et al. 2013). However, in soy bean it was shown that even a slight modification of the motif in GmFT1a (pPGWRQ) is sufficient to determine its function as a floral repressor (Liu et al. 2018). More generally, a repressing function was shown to be associated with a severe alteration or absence of the YAPGWRQ motif (Pin et al. 2010; Navarro et al. 2011; Harig et al. 2012; Lee et al. 2013; Manoharan et al. 2016; Wang et al. 2019; Yang et al. 2019).

Another well studied motif is the LYN triad, which is described to be essential for an inducing FT function (Hanzawa et al. 2005; Ahn et al. 2006; Ho and Weigel 2014). Besides containing the YAPGWRQ motif, DpFT1, DpFT2, DpFT3, DpFT5 are also the only DpFTs that contain the LYN triad, indicating that these four FTs could be potential candidates for inducing FTs. The other DpFTs show alterations in both the YAPGWRQ motif, as well as the LYN triad, directing towards a potentially repressing function. In DpFT5, the REY motif, which is the third well-studied FT motif, the tyrosine is substituted by a histamine, resulting in REH at this position. While the Y residue is described to be conserved in FTs, a H is conserved in TFLs. Swapping these amino acids with each other reversed the roles of FT and TFL in *A. thaliana* (Hanzawa et al. 2005). However, FTs from banana which contain a REH motif were able to rescue an *ft-10* mutant in Arabidopsis (Chaurasia et al. 2017), indicating that the REY motif might not be entirely necessary for an inducing function.

### *DpFT* expression in Chinese yam tubers

Based on the previously published tuber transcriptome (Riekötter et al. 2023), we analyzed the expression pattern of all DpPEBPs. Of the 75 identified PEBPs in this study, 21 FTs and 14 TFLs and 3 MFTs were expressed in the tuber tissue, underlining the diverse roles of PEBPs in growth and development in plants (Carmel-Goren et al. 2003; Chardon and Damerval 2005; McGarry and Ayre 2012; Zheng et al. 2016; Jin et al. 2021; Liu et al. 2021; Tribble et al. 2021). For most *DpPEBPs* expressed in tuber tissue, the highest expression was found in the tuber head. This is interesting, because the yam tuber head is assumed to enter a dormant stage early on, while the tip is still growing, and is assumed to be metabolically less active (Orkwor et al. 1998; Ile et al. 2006; Kawasaki et al. 2008). *DpFTs* with detectable expression in tubers showed the highest transcript levels in the head or middle part of the tubers. Similar to the known positive regulatory feed-back loop described for StSP6A in potato tubers, elevated transcript levels in the upper tuber parts might be caused by longer transcriptional activity compared to the tuber tip (Navarro et al. 2011). *DpFT7a-c* variants showed the highest expression of all *DpPEBPs* in tuber tissue but was excluded from further analysis here, because it does not contain the amino acids described to be necessary for an activating function. However, it will be interesting to investigate its function in yam tubers in future studies.

### DpFT3 is a candidate gene for a yam tuber-inducing factor

DpFT3a was investigated as a potential tuber-inducing FT (‘tuberigen’) because it was expressed in the tuber of the existing transcriptome that contained all motifs described to be necessary for an inducing function. DpFT3a localized to the nucleus in *N. benthamiana* mesophyll cells, which is in accordance with the nuclear localization described for other FTs (Abe et al. 2005; Liu et al. 2018; Kim et al. 2022). *In planta*, interaction between DpFT3a and DpFD1 was shown in *N. benthamiana* mesophyll cells indicating that DpFT3a fulfils the requirement to form a potential TAC on protein level (Abe et al. 2005; Harig et al. 2012; Teo et al. 2017; Beinecke et al. 2018; Zhang et al. 2020). Upon overexpression of *DpFT3* in *A. thaliana*, transgenic lines flowered significantly earlier than WT, indicating a flower promoting function of DpFT3 in Arabidopsis (Navarro et al. 2011; Chaurasia et al. 2017; Yang et al. 2019). This is in agreement with the previous observation that StSP6A can induce flowering in the non-flowering Arabidopsis *ft-1* mutant (Navarro et al. 2011). Interestingly, overexpression of *DpFT3a* in potato induced a strong, dosage-dependent, tuberous-storage-organ-inducing phenotype. At the same time, the overexpression did not induce flowers, indicating that DpFT3a can function as a tuber-inducing FT. Although StSP3D has been first described to play an important role in flowering as florigen, only recently a tuber-inducing activity was suggested for StSP3D in potato (Jing et al. 2023). Thus, further investigations will be required to show if DpFT3a also has a dual activity as florigen and tuberigen. Our results furthermore indicate that storage organ promoting FTs depend on other geophyte-specific factors (not present in *Arabidopsis*) to reveal their storage organ inducing function. The strong tuber-inducing phenotype of constitutively overexpressing *DpFT3a* potatoes was characterized by the absence of leaves and roots. All present shoots had turned into tuberous organs. This indicates that DpFT3a potentially has a strong effect on meristem identity, plant architecture and /or sugar flux, as was shown for other PEBP proteins. In accordance with our results, members of the PEBP-family, especially TFLs and the FT:TFL ratio, are described to play an important role in plant architecture (Carmel-Goren et al. 2003; McGarry and Ayre 2012; Jin et al. 2021; Liu et al. 2021). A regulatory role in assimilate allocation from source to sink organs has been proposed for StSP6A as leaf/stem-specific overexpression of StSP6A resulted in more efficient transport of esculin, a sucrose analogue, into the phloem for long-distance transport compared to the WT control (Lehretz et al. 2019; Lehretz et al. 2021). Furthermore, these transgenic lines revealed a reduced shoot growth due to the inhibition of stem elongation and secondary growth as well as repressed bud outgrowth as a proposed consequence of low sucrose levels, while tuber number was elevated (Lehretz et al. 2019; Lehretz et al. 2021). Therefore, DpFT3a could possess a similar function in sugar transport: inhibiting sucrose efflux from phloem to parenchyma cells and increasing the long-distance transport resulting in reduced shoot growth. Elevated sugar levels in the sink tissue might have promoted tuber development, which then served as storage organ of the newly synthesized starch that was detectable in the TC-lines. Taken together, our results strongly suggest that DpFT3 has tuber-inducing activity in geophytic plants.

## Abbreviations

FT: FLOWERING LOCUS T
SP6A: SELF PRUNING 6A
St: *Solanum tuberosum*
At: *Arabidopsis thaliana*
Dp: *Dioscorea polystachya*
FD: FLOWERING LOCUS D
TFL1: Terminal Flower 1
TSF: Twin Sister of FT
MFT: Mother of FT and TFL
BFT: Brother of FT
BiFC: Bimolecular Fluorescence Complementation
SAM: Shoot apical meristem
IM: Inflorescence meristem
CO: CONSTANS
PEBP: Phosphatidylethanolamine-binding protein
SD: Short day
LD: Long day
SP5G: SELF PRUNING 5G
TAC: Tuber Activation Complex
WT: wild type

## Acknowledgements

The authors would like to thank Verena Blome (Faculty of Chemistry and Chemical Biology, Technical University of Dortmund, Germany, former member of the Institute of Plant Biology and Biotechnology, Münster, Germany), Katrin Schrödter and Sascha Ahrens (Institute of Plant Biology and Biotechnology, Münster, Germany) for technical assistance, Dirk Prüfer (Institute of Plant Biology and Biotechnology, Münster, Germany; Fraunhofer Institute of Molecular Biology and Applied Ecology (IME), Münster, Germany) for ideas on experimental designs and supervision. Further thanks go to Theodore Drell for proofreading of the manuscript.

## Statements and Declarations

## Funding

This work was partially supported by the German Federal Ministry of Education and Research (grant number 031B0202).

## Competing Interests

The authors declare that the research was conducted in the absence of any commercial or financial relationships that could be construed as a potential conflict of interest.

## Author Contributions

Tatjana Ried and Janina Epping designed the study. Tatjana Ried conducted all experiments except the genome assembly, which was conducted by Marie Bolger and the mapping of RNAseq reads against the genome, which was conducted by Toshiyuki Sakai. Jenny Riekötter contributed to the data analysis and figure design. The manuscript was written by Tatjana Ried, Janina Epping, Jenny Riekötter and Marie Bolger. All authors read and approved the final manuscript.

## Data Availability

The datasets generated during and/or analyzed during the current study are available at the European Nucleotide Archive (ENA), Project PRJEB82723 (https://www.ebi.ac.uk/ena/browser/view/PRJEB82723).

## Notes

### Competing Interest Statement

The authors have declared no competing interest.

## References

Abe M, Kobayashi Y, Yamamoto S, Daimon Y, Yamaguchi A, Ikeda Y, Ichinoki H, Notaguchi M, Goto K, Araki T (2005) FD, a bZIP protein mediating signals from the floral pathway integrator FT at the shoot apex. Science (80-) 309:1052–1056. 10.1126/science.1115983

Abelenda JA, Cruz-Oró E, Franco-Zorrilla JM, Prat S (2016) Potato StCONSTANS-like1 Suppresses Storage Organ Formation by Directly Activating the FT-like StSP5G Repressor. Curr Biol 26:872–881. 10.1016/j.cub.2016.01.066

Ahn JH, Miller D, Winter VJ, Banfield MJ, Lee JH, Yoo SY, Henz SR, Brady RL, Weigel D (2006) A divergent external loop confers antagonistic activity on floral regulators FT and TFL1. EMBO J 25:605–614. 10.1038/SJ.EMBOJ.7600950

An H, Roussot C, Suárez-López P, Corbesier L, Vincent C, Piñeiro M, Hepworth S, Mouradov A, Justin S, Turnbull C, Coupland G (2004) CONSTANS acts in the phloem to regulate a systemic signal that induces photoperiodic flowering of Arabidopsis. Development 131:3615–3626. 10.1242/dev.01231

Arnau G, Nemorin A, Maledon E, Abraham K (2009) Revision of ploidy status of Dioscorea alata L. (Dioscoreaceae) by cytogenetic and microsatellite segregation analysis. Theor Appl Genet 118:1239–1249. 10.1007/s00122-009-0977-6

Asiedu R, Wanyera NM, Ng SYC, Ng NQ (1997) Yams. Biodiversity in trust: conservation and use of plant genetic resources in CGIAR centres. Cambridge Univ Press 57–66

Ayensu ES, Coursey DG (1972) Guinea yams the botany, ethnobotany, use and possible future of yams in West Africa. Econ Bot 26:301–318. 10.1007/BF02860700

Babil P, Ondo SK, Wata HI, Ushikawa SK, Hiwachi HS (2013) Intra-Specific Ploidy Variations in Cultivated Chinese Yam (Dioscorea polystachya Turcz.). Trop Agric Dev 57:101–107. 10.11248/jsta.57.101

Beinecke FA, Grundmann L, Wiedmann DR, Schmidt FJ, Caesar AS, Zimmermann M, Lahme M, Twyman RM, Prüfer D, Noll GA (2018) The FT/FD-dependent initiation of flowering under long-day conditions in the day-neutral species Nicotiana tabacum originates from the facultative short-day ancestor Nicotiana tomentosiformis. Plant J 96:329–342. 10.1111/tpj.14033

Bennett T, Dixon LE (2021) Asymmetric expansions of FT and TFL1 lineages characterize differential evolution of the EuPEBP family in the major angiosperm lineages. BMC Biol 19. 10.1186/S12915-021-01128-8

Bhattacharjee R, Gedil M, Sartie AM, Otoo E, Dumet D, Kikuno H, Kumar PL, Asiedu R (2011) Chapter 4 Dioscorea. Elsevier, pp 1–62

Bolger M, Schwacke R, Usadel B (2021) MapMan Visualization of RNA-Seq Data Using Mercator4 Functional Annotations. In: Dobnik D, Gruden K, Ramšak Ž, Coll A (eds). Springer US, New York, NY, pp 195–212

Bredeson J V, Lyons JB, Oniyinde IO, Okereke NR, Kolade O, Nnabue I, Nwadili CO, Hřibová E, Parker M, Nwogha J, Shu S, Carlson J, Kariba R, Muthemba S, Knop K, Barton GJ, Sherwood A V, Lopez-Montes A, Asiedu R, Jamnadass R, Muchugi A, Goodstein D, Egesi CN, Featherston J, Asfaw A, Simpson GG, Doležel J, Hendre PS, Van Deynze A, Kumar PL, Obidiegwu JE, Bhattacharjee R, Rokhsar DS (2022) Chromosome evolution and the genetic basis of agronomically important traits in greater yam. Nat Commun 13:2001. 10.1038/s41467-022-29114-w

Burkill IH (1960) The organography and the evolution of Dioscoreaceae, the family of the Yams. J Linn Soc London, Bot 56:319–412. 10.1111/j.1095-8339.1960.tb02508.x

Carmel-Goren L, Liu YS, Lifschitz E, Zamir D (2003) The self-pruning gene family in tomato. Plant Mol Biol 52:1215–1222. 10.1023/B:PLAN.0000004333.96451.11

Chardon F, Damerval C (2005) Phylogenomic Analysis of the PEBP Gene Family in Cereals. J Mol Evol 61:579–590. 10.1007/s00239-004-0179-4

Chaurasia AK, Patil HB, Krishna B, Subramaniam VR, Sane P V, Sane AP (2017) Flowering time in banana (Musa spp.), a day neutral plant, is controlled by at least three FLOWERING LOCUS T homologues. Sci Rep 7:1–14. 10.1038/s41598-017-06118-x

Chen C, Chen H, Zhang Y, Thomas HR, Frank MH, He Y, Xia R (2020) TBtools: An Integrative Toolkit Developed for Interactive Analyses of Big Biological Data. Mol Plant 13:1194–1202. 10.1016/j.molp.2020.06.009

Chen S, Zhou Y, Chen Y, Gu J (2018) Fastp: An ultra-fast all-in-one FASTQ preprocessor. Bioinformatics 34:i884– i890. 10.1093/bioinformatics/bty560

Cheng H, Concepcion GT, Feng X, Zhang H, Li H (2021) Haplotype-resolved de novo assembly using phased assembly graphs with hifiasm. Nat Methods 18:170–175. 10.1038/s41592-020-01056-5

Corbesier L, Vincent C, Jang S, Fornara F, Fan Q, Searle I, Giakountis A, Farrona S, Gissot L, Turnbull C, Coupland G (2007) FT protein movement contributes to long-distance signaling in floral induction of Arabidopsis. Science (80-) 316:1030–1033. 10.1126/science.1141752

Coursey DG (1967) Yams - An account of the Nature, Origins, Cultivation and Utilisation of the Useful Members of the Dioscoreaceae. Longmans, Green Co LTD, London

Dong L, Lu Y, Liu S (2020) Genome-wide member identification, phylogeny and expression analysis of PEBP gene family in wheat and its progenitors. PeerJ 8:1–22. 10.7717/peerj.10483

Fan Y, He Q, Luo A, Wang M, Luo A (2015) Characterization and antihyperglycemic activity of a polysaccharide from Dioscorea opposita Thunb roots. Int J Mol Sci 16:6391–6401. 10.3390/ijms16036391

Gao X, Li B, Jiang H, Liu F, Xu D, Liu Z (2007) Dioscorea opposita reverses dexamethasone induced insulin resistance. Fitoterapia 78:12–15. 10.1016/j.fitote.2006.09.015

González-Schain ND, Díaz-Mendoza M, Zurczak M, Suárez-López P (2012) Potato CONSTANS is involved in photoperiodic tuberization in a graft-transmissible manner. Plant J 70:678–690. 10.1111/j.1365-313X.2012.04909.x

Hahn, S. K. Osiru, D.S.O. Akoroda, M.O. and Otoo JA (1987). (1987) Yam production and its future prospects. Outlook on Agriculture,. Agriculture, 16:105 110.

Hanzawa Y, Money T, Bradley D (2005) A single amino acid converts a repressor to an activator of flowering. Proc Natl Acad Sci U S A 102:7748–7753. 10.1073/pnas.0500932102

Harig L, Beinecke FA, Oltmanns J, Muth J, Müller O, Rüping B, Twyman RM, Fischer R, Prüfer D, Noll GA (2012) Proteins from the FLOWERING LOCUS T-like subclade of the PEBP family act antagonistically to regulate floral initiation in tobacco. Plant J 72:908–921. 10.1111/j.1365-313X.2012.05125.x

Ho WWH, Weigel D (2014) Structural features determining flower-promoting activity of Arabidopsis FLOWERING LOCUS T. Plant Cell 26:552–564. 10.1105/tpc.113.115220

Hoekema A, Hirsch PR, Hooykaas PJJ, Schilperoort RA (1983) A binary plant vector strategy based on separation of vir- and T-region of the Agrobacterium tumefaciens Ti-plasmid. Nature 303:179–180. 10.1038/303179a0

Holst F, Bolger A, Günther C, Maß J, Triesch S, Kindel F, Kiel N, Saadat N, Ebenhöh O, Usadel B, Schwacke R, Bolger M, Weber APM, Denton AK (2023) Helixer–de novo Prediction of Primary Eukaryotic Gene Models Combining Deep Learning and a Hidden Markov Model. bioRxiv 10.1101/2023.02.06.527280.

Horsch RB, Fry JE, Hoffmann NL, Eichholtz D, Rogers SG, Fraley RT (1985) A simple and general method for transferring genes into plants. Science 227:1229–1231. 10.1126/science.227.4691.1229

Ile EI, Craufurd PQ, Battey NH, Asiedu R (2006) Phases of dormancy in yam tubers (Dioscorea rotundata). Ann Bot 97:497–504. 10.1093/aob/mcl002

Irvine FR (1952) Supplementary and emergency food plants of West Africa. Econ Bot 6:23–40. 10.1007/BF02859192

Jin S, Nasim Z, Susila H, Ahn JH (2021) Evolution and functional diversification of FLOWERING LOCUS T/TERMINAL FLOWER 1 family genes in plants. Semin Cell Dev Biol 109:20–30. 10.1016/j.semcdb.2020.05.007

Jing S, Jiang P, Sun X, Yu L, Wang E, Qin J, Zhang F, Prat S, Song B (2023) Long-distance control of potato storage organ formation by SELF PRUNING 3D and FLOWERING LOCUS T-like 1. Plant Commun 4:100547. 10.1016/j.xplc.2023.100547

Kardailsky I, Shukla VK, Ahn JH, Dagenais N, Christensen SK, Nguyen JT, Chory J, Harrison MJ, Weigel D (1999) Activation tagging of the floral inducer FT. Science (80-) 286:1962–1965. 10.1126/science.286.5446.1962

Karlgren A, Gyllenstrand N, Källman T, Sundström JF, Moore D, Lascoux M, Lagercrantz U (2011) Evolution of the PEBP gene family in plants: Functional diversification in seed plant evolution. Plant Physiol 156:1967–1977. 10.1104/pp.111.176206

Kawasaki M, Taniguchi M, Miyake H (2008) Dynamics of amyloplast sedimentation in growing yam tubers and its possible role in graviperception. Plant Prod Sci 11:393–397. 10.1626/pps.11.393

Khosa J, Bellinazzo F, Kamenetsky Goldstein R, Macknight R, Immink RGH (2021) PHOSPHATIDYLETHANOLAMINE-BINDING PROTEINS: The conductors of dual reproduction in plants with vegetative storage organs. J Exp Bot 72:2845–2856. 10.1093/jxb/erab064

Kim D, Langmead B, Salzberg SL (2015) HISAT: A fast spliced aligner with low memory requirements. Nat Methods 12:357–360. 10.1038/nmeth.3317

Kim G, Rim Y, Cho H, Hyun TK (2022) Identification and Functional Characterization of FLOWERING LOCUS T in Platycodon grandiflorus. Plants 11:325. 10.3390/PLANTS11030325/S1

Kobayashi Y (1999) A Pair of Related Genes with Antagonistic Roles in Mediating Flowering Signals. Science (80-) 286:1960–1962. 10.1126/science.286.5446.1960

Koornneef M, Hanhart CJ, van der Veen JH (1991) A genetic and physiological analysis of late flowering mutants in Arabidopsis thaliana. Mol Gen Genet 229:57–66. 10.1007/BF00264213

Kwon C-S, Sohn HY, Kim SH, Kim JH, Son KH, Lee JS, Lim JK, Kim J-S (2003) Anti-obesity Effect of Dioscorea nipponica Makino with Lipase-inhibitory Activity in Rodents. Biosci Biotechnol Biochem 67:1451– 1456. 10.1271/bbb.67.1451

Lee J, Oh M, Park H, Lee I (2008) SOC1 translocated to the nucleus by interaction with AGL24 directly regulates LEAFY. Plant J 55:832–843. 10.1111/j.1365-313X.2008.03552.x

Lee R, Baldwin S, Kenel F, McCallum J, Macknight R (2013) FLOWERING LOCUS T genes control onion bulb formation and flowering. Nat Commun 4:1–9. 10.1038/ncomms3884

Lehretz GG, Sonnewald S, Hornyik C, Corral JM, Sonnewald U (2019) Post-transcriptional Regulation of FLOWERING LOCUS T Modulates Heat-Dependent Source-Sink Development in Potato. Curr Biol 29:1614– 1624. 10.1016/j.cub.2019.04.027

Lehretz GG, Sonnewald S, Sonnewald U (2021) Assimilate highway to sink organs – Physiological consequences of SP6A overexpression in transgenic potato (Solanum tuberosum L.). J Plant Physiol 266:153530. 10.1016/j.jplph.2021.153530

Liao Y, Smyth GK, Shi W (2014) featureCounts: an efficient general purpose program for assigning sequence reads to genomic features. Bioinformatics 30:923–930. 10.1093/bioinformatics/btt656

Lin MK, Belanger H, Lee YJ, Varkonyi-Gasic E, Taoka KI, Miura E, Xoconostle-Cázares B, Gendler K, Jorgensen RA, Phinney B, Lough TJ, Lucas WJ (2007) FLOWERING LOCUS T protein may act as the long-distance florigenic signal in the cucurbits. Plant Cell 19:1488–1506. 10.1105/tpc.107.051920

Liu H, Huang X, Ma B, Zhang T, Sang N, Zhuo L, Zhu J (2021) Components and Functional Diversification of Florigen Activation Complexes in Cotton. Plant Cell Physiol 62:1542–1555. 10.1093/pcp/pcab107

Liu W, Jiang B, Ma L, Zhang S, Zhai H, Xu X, Hou W, Xia Z, Wu C, Sun S, Wu T, Chen L, Han T (2018) Functional diversification of Flowering Locus T homologs in soybean: GmFT1a and GmFT2a/5a have opposite roles in controlling flowering and maturation. New Phytol 217:1335–1345. 10.1111/nph.14884

Liu Y-H, Lin Y-S, Liu D-Z, Han C-H, Chen C-T, Fan M, Hou W-C (2009) Effects of Different Types of Yam (Dioscorea alata) Products on the Blood Pressure of Spontaneously Hypertensive Rats. Biosci Biotechnol Biochem 73:1371–1376. 10.1271/bbb.90022

Livak KJ, Schmittgen TD (2001) Analysis of relative gene expression data using real-time quantitative PCR and the 2(−Delta Delta C(T)) Method. Methods 25:402–408. 10.1006/meth.2001.1262

Mabhaudhi T, Chimonyo VGP, Hlahla S, Massawe F, Mayes S, Nhamo L, Modi AT (2019) Prospects of orphan crops in climate change. Planta 250:695–708. 10.1007/s00425-019-03129-y

Manni M, Berkeley MR, Seppey M, Simão FA, Zdobnov EM (2021) BUSCO Update: Novel and Streamlined Workflows along with Broader and Deeper Phylogenetic Coverage for Scoring of Eukaryotic, Prokaryotic, and Viral Genomes. Mol Biol Evol 38:4647–4654. 10.1093/molbev/msab199

Manoharan RK, Han JSH, Vijayakumar H, Subramani B, Thamilarasan SK, Park JI, Nou IS (2016) Molecular and functional characterization of FLOWERING LOCUS T homologs in Allium cepa. Molecules 21:1–14. 10.3390/molecules21020217

Martinez-Garcia JF, Virgos-Soler A, Prat S (2002) Control of photoperiod-regulated tuberization in potato by the Arabidopsis flowering-time gene CONSTANS. Proc Natl Acad Sci 99:15211–15216. 10.1073/pnas.222390599

McGarry RC, Ayre BG (2012) Manipulating plant architecture with members of the CETS gene family. Plant Sci 188– 189:71–81. 10.1016/j.plantsci.2012.03.002

Mignouna HD, Abang MM, Asiedu R (2007) Advances in yam (Dioscorea spp.) genetics and genomics. Proc 13th ISTRC Symp 72–81

Mignouna HD, Abang MM, Asiedu R (2003) Harnessing modern biotechnology for tropical tuber crop improvement: Yam (Dioscorea spp.) molecular breeding. African J Biotechnol 2:478–492

Navarro C, Abelenda JA, Cruz-Oró E, Cuéllar CA, Tamaki S, Silva J, Shimamoto K, Prat S (2011) Control of flowering and storage organ formation in potato by FLOWERING LOCUS T. Nature 478:119–122. 10.1038/nature10431

O’Sullivan J. N. (2010) Yam nutrition: nutrient disorders and soil fertility management. Australian Centre for International Agricultural Research (ACIAR), Canberra

Opara LU (2003) YAMS: Post-harvest Operations. INPhO - Post-harvest Compend 23

Orkwor GC, Asiedu R, Ekanayaka IJ (1998) Food Yams - Advances in Research

Otoo E (2017) Yam Breeding in Ghana. J Agric Sci 9:122. 10.5539/jas.v9n10p122

Pin PA, Benlloch R, Bonnet D, Wremerth-Weich E, Kraft T, Gielen JJL, Nilsson O (2010) An antagonistic pair of FT homologs mediates the control of flowering time in sugar beet. Science 330:1397–400. 10.1126/science.1197004

Riekötter J, Oklestkova J, Muth J, Twyman RM, Epping J (2023) Transcriptomic analysis of Chinese yam (Dioscorea polystachya Turcz.) variants indicates brassinosteroid involvement in tuber development. Front Nutr 1–20. 10.3389/fnut.2023.1112793

Schwacke R, Ponce-Soto GY, Krause K, Bolger AM, Arsova B, Hallab A, Gruden K, Stitt M, Bolger ME, Usadel B (2019) MapMan4: A Refined Protein Classification and Annotation Framework Applicable to Multi-Omics Data Analysis. Mol Plant 12:879–892. 10.1016/j.molp.2019.01.003

Scott G, Best R, Rosegrant M, Bokanga M (2000) Root and tuber in the global food system: a vision statement to the year 2020. Lima, Peru: International Potato Center

Shannon S, Meeks-Wagner DR (1993) Genetic Interactions That Regulate Inflorescence Development in Arabidopsis. Plant Cell 5:639–655. 10.1105/tpc.5.6.639

Shannon S, Meeks-Wagner DR (1991) A Mutation in the Arabidopsis TFL1 Gene Affects Inflorescence Meristem Development. Plant Cell 3:877–892. 10.1105/TPC.3.9.877

Sharma P, Lin T, Hannapel DJ (2016) Targets of the StBEL5 transcription factor include the FT Ortholog StSP6A1[OPEN]. Plant Physiol 170:310–324. 10.1104/pp.15.01314

Shewry PR (2003) Tuber storage proteins. Ann Bot 91:755–769. 10.1093/aob/mcg084

Simão FA, Waterhouse RM, Ioannidis P, Kriventseva E V., Zdobnov EM (2015) BUSCO: Assessing genome assembly and annotation completeness with single-copy orthologs. Bioinformatics 31:3210–3212. 10.1093/bioinformatics/btv351

Son IS, Lee JS, Lee JY, Kwon CS (2014) Antioxidant and Anti-inflammatory Effects of Yam (Dioscorea batatas Decne.) on Azoxymethane-induced Colonic Aberrant Crypt Foci in F344 Rats. Prev Nutr food Sci 19:82–88. 10.3746/pnf.2014.19.2.082

Stolze A, Wanke A, van Deenen N, Geyer R, Prüfer D, Schulze Gronover C (2017) Development of rubber-enriched dandelion varieties by metabolic engineering of the inulin pathway. Plant Biotechnol J 15:740–753. s10.1111/pbi.12672

Susila H, Purwestri YA (2023) PEBP Signaling Network in Tubers and Tuberous Root Crops. Plants 12:1–15. 10.3390/plants12020264

Tamaki S, Matsuo S, Wong HL, Yokoi S, Shimamoto K (2007) Hd3a Protein Is a Mobile Flowering Signal in Rice. Science (80-) 316

Tamiru M, Natsume S, Takagi H, White B, Yaegashi H, Shimizu M, Yoshida K, Uemura A, Oikawa K, Abe A, Urasaki N, Matsumura H, Babil P, Yamanaka S, Matsumoto R, Muranaka S, Girma G, Lopez-Montes A, Gedil M, Bhattacharjee R, Abberton M, Kumar PL, Rabbi I, Tsujimura M, Terachi T, Haerty W, Corpas M, Kamoun S, Kahl G, Takagi H, Asiedu R, Terauchi R (2017) Genome sequencing of the staple food crop white Guinea yam enables the development of a molecular marker for sex determination. BMC Biol 15:1–20. 10.1186/s12915-017-0419-x

Tamura K, Stecher G, Kumar S (2021) MEGA11: Molecular Evolutionary Genetics Analysis Version 11. Mol Biol Evol 38:3022–3027. 10.1093/molbev/msab120

Taoka K, Ohki I, Tsuji H, Furuita K, Hayashi K, Yanase T, Yamaguchi M, Nakashima C, Purwestri YA, Tamaki S, Ogaki Y, Shimada C, Nakagawa A, Kojima C, Shimamoto K (2011) 14-3-3 proteins act as intracellular receptors for rice Hd3a florigen. Nature 476:332–335. 10.1038/nature10272

Teo CJ, Takahashi K, Shimizu K, Shimamoto K, Taoka KI (2017) Potato tuber induction is regulated by interactions between components of a tuberigen complex. Plant Cell Physiol 58:365–374. 10.1093/pcp/pcw197

Tribble CM, Martínez-Gómez J, Alzate-Guarín F, Rothfels CJ, Specht CD (2021) Comparative transcriptomics of a monocotyledonous geophyte reveals shared molecular mechanisms of underground storage organ formation ***

Tribble, C.M. et al. (2021) ‘Comparative transcriptomics of a monocotyledonous geophyte reveals shared molecular mechanis. Evol Dev. 10.1111/ede.12369

Turck F, Fornara F, Coupland G (2008) Regulation and Identity of Florigen: FLOWERING LOCUS T Moves Center Stage. Annu Rev Plant Biol 59:573–594. 10.1146/annurev.arplant.59.032607.092755

Wang L, Yan J, Zhou X, Cheng S, Chen Z, Song Q, Liu X, Ye J, Zhang W, Wu G, Xu F (2019) GbFT, a FLOWERING LOCUS T homolog from Ginkgo biloba, promotes flowering in transgenic Arabidopsis. Sci Hortic (Amsterdam) 247:205–215. 10.1016/j.scienta.2018.12.020

Waterhouse AM, Procter JB, Martin DMA, Clamp M, Barton GJ (2009) Jalview Version 2-A multiple sequence alignment editor and analysis workbench. Bioinformatics 25:1189–1191. 10.1093/bioinformatics/btp033

Wigge PA, Kim MC, Jaeger KE, Busch W, Schmid M, Lohmann JU, Weigel D (2005) Integration of spatial and temporal information during floral induction in Arabidopsis. Science (80-) 309:1056–1059. 10.1126/science.1114358

Xing S, van Deenen N, Magliano P, Frahm L, Forestier E, Nawrath C, Schaller H, Gronover CS, Prüfer D, Poirier Y (2014) ATP citrate lyase activity is post-translationally regulated by sink strength and impacts the wax, cutin and rubber biosynthetic pathways. Plant J 79:270–284. 10.1111/tpj.12559

Xu X, Pan S, Cheng S, Zhang B, Mu D, Ni P, Zhang G, Yang S, Li R, Wang J, Orjeda G, Guzman F, Torres M, Lozano R, Ponce O, Martinez D, De la Cruz G, Chakrabarti SK, Patil VU, Skryabin KG, Kuznetsov BB, Ravin N V., Kolganova T V., Beletsky A V., Mardanov A V., Di Genova A, Bolser DM, Martin DMA, Li G, Yang Y, Kuang H, Hu Q, Xiong X, Bishop GJ, Sagredo B, Mejía N, Zagorski W, Gromadka R, Gawor J, Szczesny P, Huang S, Zhang Z, Liang C, He J, Li Y, He Y, Xu J, Zhang Y, Xie B, Du Y, Qu D, Bonierbale M, Ghislain M, del Rosario Herrera M, Giuliano G, Pietrella M, Perrotta G, Facella P, O’Brien K, Feingold SE, Barreiro LE, Massa GA, Diambra L, Whitty BR, Vaillancourt B, Lin H, Massa AN, Geoffroy M, Lundback S, DellaPenna D, Robin Buell C, Sharma SK, Marshall DF, Waugh R, Bryan GJ, Destefanis M, Nagy I, Milbourne D, Thomson SJ, Fiers M, Jacobs JME, Nielsen KL, Sønderkær M, Iovene M, Torres GA, Jiang J, Veilleux RE, Bachem CWB, de Boer J, Borm T, Kloosterman B, van Eck H, Datema E, te Lintel Hekkert B, Goverse A, van Ham RCHJ, Visser RGF (2011) Genome sequence and analysis of the tuber crop potato. Nature 475:189–195. 10.1038/nature10158

Yang Z, Chen L, Kohnen M V, Xiong B, Zhen X, Liao J, Oka Y, Zhu Q, Gu L, Lin C, Liu B (2019) Identification and Characterization of the PEBP Family Genes in Moso Bamboo (Phyllostachys heterocycla). Sci Rep 9:1–12. 10.1038/s41598-019-51278-7

Yoo SC, Chen C, Rojas M, Daimon Y, Ham BK, Araki T, Lucas WJ (2013) Phloem long-distance delivery of FLOWERING LOCUS T (FT) to the apex. Plant J 75:456–468. 10.1111/tpj.12213

Zhang M, Li P, Yan X, Wang J, Cheng T, Zhang Q (2021) Genome-wide characterization of PEBP family genes in nine Rosaceae tree species and their expression analysis in P. mume. BMC Ecol Evol 21:1–23. 10.1186/s12862-021-01762-4

Zhang X, Campbell R, Ducreux LJM, Morris J, Hedley PE, Mellado-Ortega E, Roberts AG, Stephens J, Bryan GJ, Torrance L, Chapman SN, Prat S, Taylor MA (2020) TERMINAL FLOWER-1/CENTRORADIALIS inhibits tuberisation via protein interaction with the tuberigen activation complex. Plant J 1–16. 10.1111/tpj.14898

Zheng XM, Wu FQ, Zhang X, Lin QB, Wang J, Guo XP, Lei CL, Cheng ZJ, Zou C, Wan JM (2016) Evolution of the PEBP gene family and selective signature on FT-like clade. J Syst Evol 54:502–510. 10.1111/jse.12199

