## Supplementary Material for "Genome Sequencing of Chinese yam (*Dioscorea polystachya*): analysis of PEBP gene family diversity and identification of a potential tuber inducing factor"

[1]Cologne, Germany

**Table S1: DpPEBP GeneID and Annotation**

| Gene_ID_1 | DpPEBP Name | Mercator description | prot-scriber | swissprot |
|---|---|---|---|---|
| d_polystachya_ptg000024l_000055.1 | FT1a | florigen component *(FT) | protein flowering locus t | HEADING DATE 3A |
| d_polystachya_ptg000164l_000049.1 | FT1b | florigen component *(FT) | protein flowering locus t | HEADING DATE 3A |
| d_polystachya_ptg000166l_000157.1 | FT1c | florigen component *(FT) | protein flowering locus t | HEADING DATE 3A |
| d_polystachya_ptg000024l_000054.1 | FT2a | florigen component *(FT) | protein flowering locus t | HEADING DATE 3A |
| d_polystachya_ptg000164l_000048.1 | FT2b | florigen component *(FT) | protein flowering locus t | HEADING DATE 3A |
| d_polystachya_ptg000166l_000156.1 | FT2c | florigen component *(FT) | protein flowering locus t | HEADING DATE 3A |
| d_polystachya_ptg000008l_000149.1 | FT3a | florigen component *(FT) | protein flowering locus t | FLOWERING LOCUS T |
| d_polystachya_ptg000008l_000151.1 | FT3b | florigen component *(FT) | protein flowering locus t | FLOWERING LOCUS T |
| d_polystachya_ptg000058l_000117.1 | FT3c | florigen component *(FT) | protein flowering locus t | FLOWERING LOCUS T |
| d_polystachya_ptg000058l_000119.1 | FT3d | florigen component *(FT) | protein flowering locus t | FLOWERING LOCUS T |
| d_polystachya_ptg000079l_000508.1 | FT3e | florigen component *(FT) | protein flowering locus t | FLOWERING LOCUS T |
| d_polystachya_ptg000079l_000512.1 | FT3f | not classified | protein mother of ft and tfl homolog | FLOWERING LOCUS T |
| d_polystachya_ptg000086l_000448.1 | FT3g | florigen component *(FT) | protein flowering locus t | FLOWERING LOCUS T |
| d_polystachya_ptg000106l_000674.1 | FT3h | florigen component *(FT) | protein flowering locus t | FLOWERING LOCUS T |
| d_polystachya_ptg000106l_000676.1 | FT3i | not classified | protein flowering locus t | FLOWERING LOCUS T |
| d_polystachya_ptg000078l_000546.1 | FT4a | not classified | protein of ft and tfl | HEADING DATE 3A |
| d_polystachya_ptg000044l_000186.1 | FT4b | not classified | protein of ft and tfl | HEADING DATE 3A |
| d_polystachya_ptg000005l_000212.1 | FT4c | not classified | isoform of protein flowering locus t | HEADING DATE 3A |
| d_polystachya_ptg000031l_000159.1 | FT5a | not classified | protein flowering locus t | FLOWERING LOCUS T |
| d_polystachya_ptg000196l_000382.1 | FT5b | not classified | protein flowering locus t | FLOWERING LOCUS T |
| d_polystachya_ptg000104l_000205.1 | FT5c | florigen component *(FT) | protein flowering locus t | HEADING DATE 3A |
| d_polystachya_ptg000089l_000192.1 | FT5d | florigen component *(FT) | protein flowering locus t | HEADING DATE 3A |
| d_polystachya_ptg000012l_000528.1 | FT6a | not classified | protein of ft and tfl | RICE FOWERING LOCUS T 1 |
| d_polystachya_ptg000160l_000562.1 | FT6b | not classified | protein of ft and tfl | RICE FOWERING LOCUS T 1 |
| d_polystachya_ptg000100l_000524.1 | FT6c | not classified | protein of ft and tfl | RICE FOWERING LOCUS T 1 |
| d_polystachya_ptg000078l_000545.1 | FT7a | not classified | isoform of protein flowering locus t | TWIN SISTER of FT |
| d_polystachya_ptg000044l_000185.1 | FT7b | not classified | protein of ft and tfl | FLOWERING LOCUS T |
| d_polystachya_ptg000005l_000210.1 | FT7c | not classified | isoform of protein flowering locus t | TWIN SISTER of FT |
| d_polystachya_ptg000063l_000722.1 | FT8a | not classified | protein flowering locus t | TWIN SISTER of FT |
| d_polystachya_ptg000004l_000199.1 | FT8b | not classified | protein flowering locus t | TWIN SISTER of FT |
| d_polystachya_ptg000093l_000706.1 | FT8c | not classified | protein flowering locus t | TWIN SISTER of FT |
| d_polystachya_ptg000011l_000447.1 | FT9a | not classified | protein flowering locus t | FLOWERING LOCUS T |

| | | | | |
|---|---|---|---|---|
| d_polystachya_ptg000010l_000250.1 | FT9b | not classified | protein flowering locus t | FLOWERING LOCUS T |
| d_polystachya_ptg000074l_000481.1 | FT9c | not classified | protein flowering locus t | FLOWERING LOCUS T |
| d_polystachya_ptg000024l_000239.1 | MFT1a | not classified | protein mother of ft and tfl homolog | Protein MOTHER of FT and TFL1 homolog 1 |
| d_polystachya_ptg000164l_000241.1 | MFT1b | not classified | protein mother of ft and tfl homolog | Protein MOTHER of FT and TFL1 homolog 1 |
| d_polystachya_ptg000180l_000084.1 | MFT1c | not classified | protein mother of ft and tfl homolog | Protein MOTHER of FT and TFL1 homolog 1 |
| d_polystachya_ptg000003l_000697.1 | MFT2a | not classified | protein mother of ft and tfl homolog | Protein MOTHER of FT and TFL1 |
| d_polystachya_ptg000057l_000739.1 | MFT2b | not classified | protein mother of ft and tfl homolog | Protein MOTHER of FT and TFL1 homolog 1 |
| d_polystachya_ptg000066l_000779.1 | MFT2c | not classified | protein mother of ft and tfl homolog | Protein MOTHER of FT and TFL1 homolog 1 |
| d_polystachya_ptg000168l_000374.1 | MFT2d | not classified | protein mother of ft and tfl homolog | Protein MOTHER of FT and TFL1 |
| d_polystachya_ptg000168l_000373.1 | MFT2e | not classified | protein mother of ft and tfl homolog | Protein MOTHER of FT and TFL1 |
| d_polystachya_ptg000168l_000371.1 | MFT2f | not classified | protein mother of ft and tfl homolog | Protein MOTHER of FT and TFL1 homolog 1 |
| d_polystachya_ptg000181l_000335.1 | MFT2g | not classified | protein mother of ft and tfl homolog | Protein MOTHER of FT and TFL1 |
| d_polystachya_ptg000003l_000699.1 | MFT2h | not classified | protein mother of ft and tfl | Protein MOTHER of FT and TFL1 homolog 1 |
| d_polystachya_ptg000181l_000336.1 | MFT2i | not classified | protein mother of ft and tfl homolog | Protein MOTHER of FT and TFL1 |
| d_polystachya_ptg000057l_000741.1 | MFT2j | not classified | protein mother of ft and tfl homolog | Protein MOTHER of FT and TFL1 |
| d_polystachya_ptg000066l_000778.1 | MFT2k | not classified | protein mother of ft and tfl homolog | Protein MOTHER of FT and TFL1 |
| d_polystachya_ptg000181l_000334.1 | MFT2l | not classified | protein mother of ft and tfl homolog | Protein MOTHER of FT and TFL1 |
| d_polystachya_ptg000003l_000696.1 | MFT2m | not classified | protein mother of ft and tfl homolog | Protein MOTHER of FT and TFL1 homolog 1 |
| d_polystachya_ptg000168l_000372.1 | MFT2n | not classified | protein mother of ft and tfl homolog | no annotation |
| d_polystachya_ptg000057l_000738.1 | MFT2o | not classified | protein mother of ft and tfl homolog | Protein MOTHER of FT and TFL1 |
| d_polystachya_ptg000066l_000780.1 | MFT2p | not classified | protein mother of ft and tfl | no annotation |
| d_polystachya_ptg000003l_000694.1 | MFT2q | not classified | protein mother of ft and tfl | no annotation |
| d_polystachya_ptg000003l_000695.1 | MFT2r | not classified | protein mother of ft and tfl | no annotation |
| d_polystachya_ptg000066l_000781.1 | MFT2s | not classified | protein mother of ft and tfl homolog | Protein MOTHER of FT and TFL1 homolog 1 |
| d_polystachya_ptg000031l_000309.1 | TFL1a | effector protein *(TFL/BFT/CEN) | protein mother of ft and tfl | CEN-like protein 2 |
| d_polystachya_ptg000089l_000349.1 | TFL1b | effector protein *(TFL/BFT/CEN) | protein mother of ft and tfl | Protein SELF_PRUNING |
| d_polystachya_ptg000104l_000374.1 | TFL1c | effector protein *(TFL/BFT/CEN) | protein mother of ft and tfl | Protein SELF_PRUNING |
| d_polystachya_ptg000196l_000539.1 | TFL1d | effector protein *(TFL/BFT/CEN) | protein mother of ft and tfl | Protein SELF_PRUNING |
| d_polystachya_ptg002384l_000005.1 | TFL1e | effector protein *(TFL/BFT/CEN) | protein mother of ft and tfl | Protein SELF_PRUNING |
| d_polystachya_ptg000058l_000293.1 | TFL2a | effector protein *(TFL/BFT/CEN) | protein mother of ft and tfl homolog | CEN-like protein 2 |
| d_polystachya_ptg000070l_000492.1 | TFL2b | effector protein *(TFL/BFT/CEN) | protein mother of ft and tfl homolog | CEN-like protein 2 |
| d_polystachya_ptg000079l_000699.1 | TFL2c | effector protein *(TFL/BFT/CEN) | protein mother of ft and tfl homolog | CEN-like protein 2 |
| d_polystachya_ptg000106l_000863.1 | TFL2d | effector protein *(TFL/BFT/CEN) | protein mother of ft and tfl homolog | CEN-like protein 2 |
| d_polystachya_ptg000003l_000492.1 | TFL3a | effector protein *(TFL/BFT/CEN) | protein mother of ft and tfl | Protein SELF_PRUNING |

| | | | | |
|---|---|---|---|---|
| d_polystachya_ptg000010l_000007.1 | TFL3b | effector protein *(TFL/BFT/CEN) | protein mother of ft and tfl | Protein SELF_PRUNING |
| d_polystachya_ptg000122l_000069.1 | TFL3c | effector protein *(TFL/BFT/CEN) | protein mother of ft and tfl | Protein SELF_PRUNING |
| d_polystachya_ptg000181l_000169.1 | TFL3d | effector protein *(TFL/BFT/CEN) | protein mother of ft and tfl | Protein SELF_PRUNING |
| d_polystachya_ptg000005l_000341.1 | TFL4a | effector protein *(TFL/BFT/CEN) | protein mother of ft and tfl | CEN-like protein 2 |
| d_polystachya_ptg000078l_000662.1 | TFL4b | effector protein *(TFL/BFT/CEN) | protein mother of ft and tfl | CEN-like protein 2 |
| d_polystachya_ptg000003l_000064.1 | TFL5a | effector protein *(TFL/BFT/CEN) | protein mother of ft and tfl homolog | CEN-like protein 1 |
| d_polystachya_ptg000057l_000284.1 | TFL5b | effector protein *(TFL/BFT/CEN) | protein mother of ft and tfl homolog | CEN-like protein 1 |
| d_polystachya_ptg000066l_000262.1 | TFL5c | effector protein *(TFL/BFT/CEN) | protein mother of ft and tfl | CEN-like protein 1 |
| d_polystachya_ptg000126l_000785.1 | TFL5d | effector protein *(TFL/BFT/CEN) | protein mother of ft and tfl homolog | CEN-like protein 1 |
| d_polystacya_ptg000058l_000423.1 | FD1 | bZIP class-A transcription factor | bzip transcription factor | bZIP transcription factor 12 |

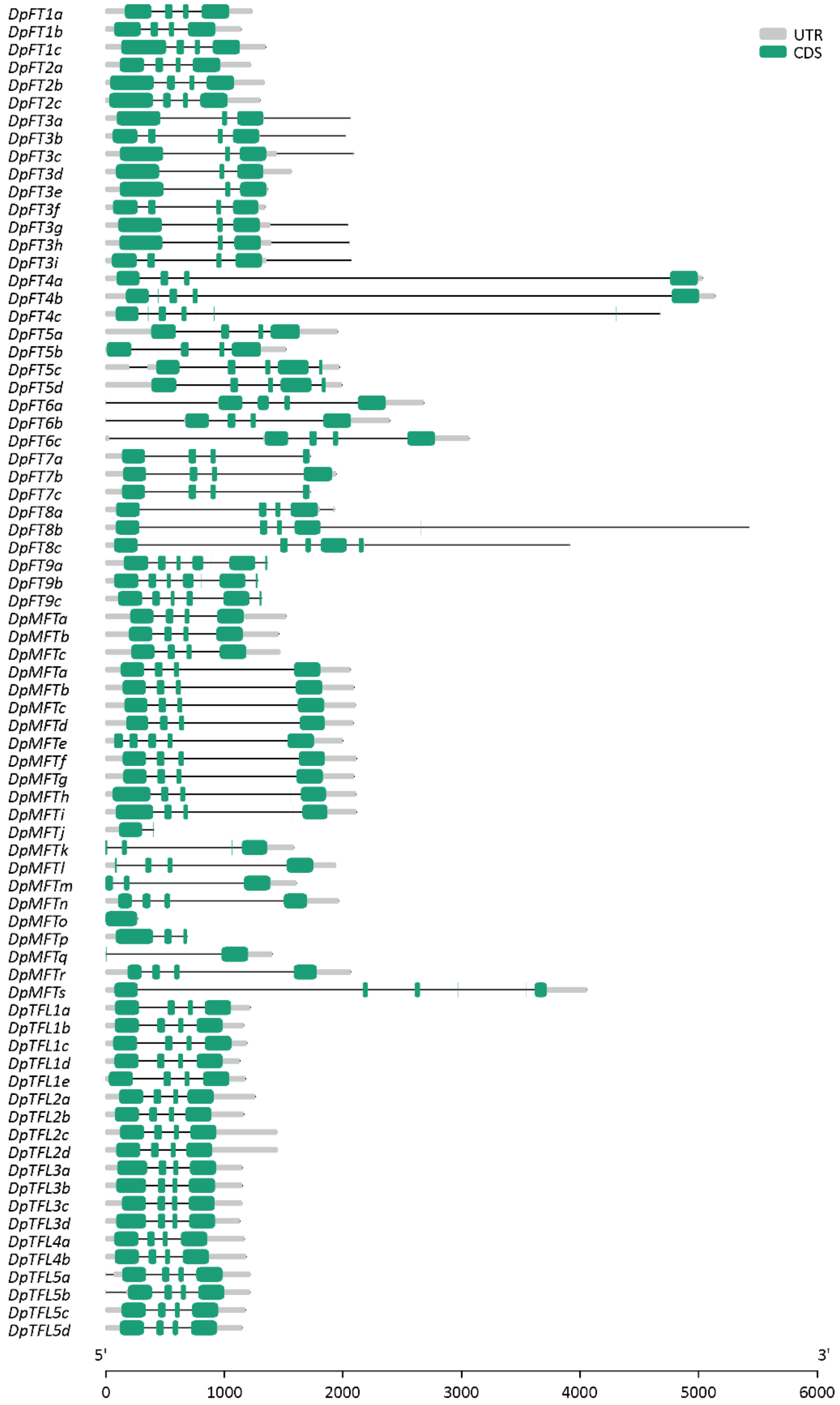


**Fig. S1.** Intron-exon structure of *DpPEBPs* Sequences. Sequences are displayed from 5’ on the left to 3’ on the right. Green boxes indicate exons, grey boxes indicate the 5’ and 3’ UTRs and black lines indicate introns. The scale bar is represented in base pairs.

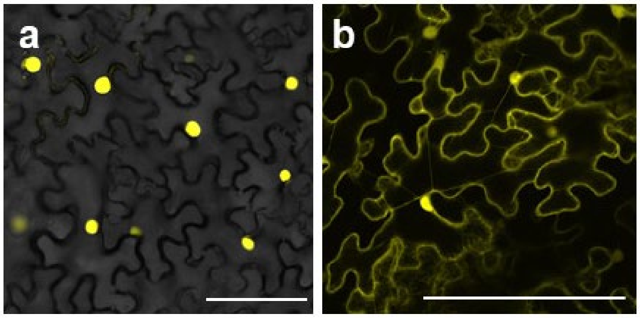


**Fig. S2.** Cellular localization of DpFD1 and DpFT3a. CLSM images of *N. benthamiana* epidermal cells transiently expressing DpFD1 and DpFT3a. For visualization of localization, the YFP variant Venus was fused to the N-terminus of DpFT3a and DpFD1. Nuclear localization of Venus-DpFD1 (a); Cytoplasmatic and nuclear localization of Venus-DpFT3a (b). Scale bar: 100 µm.

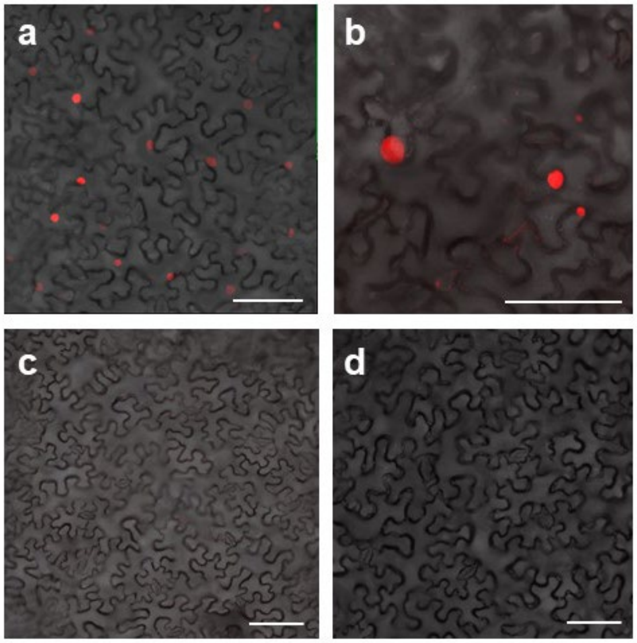


**Fig. S3.** Interaction of DpFD1 and DpFT3a. CLSM images of *N. benthamiana* epidermal cells transiently expressing DpFD1 and DpFT3a proteins. To visualize a potential interaction between DpFT3a and DpFD1, the split-mRFP system was used: The N-terminal and C-terminal regions of mRFP were fused to the N-terminus of DpFT3a and DpFD1, respectively. Detection of red fluorescence in the nucleus of epidermal cells expressing N-mRFP-DpFT3a and C-mRFP-DpFD1 (a), and N-mRFP-AtFT and C-mRFP-AtFD (positive control) (b), respectively, indicating protein-protein interaction. Single infiltrations served as negative controls. No fluorescence signal was detected in epidermal cells expressing C-mRFP-DpFD1 (c) or N-mRFP-DpFT3a (d). Scale bar: 100 µm.

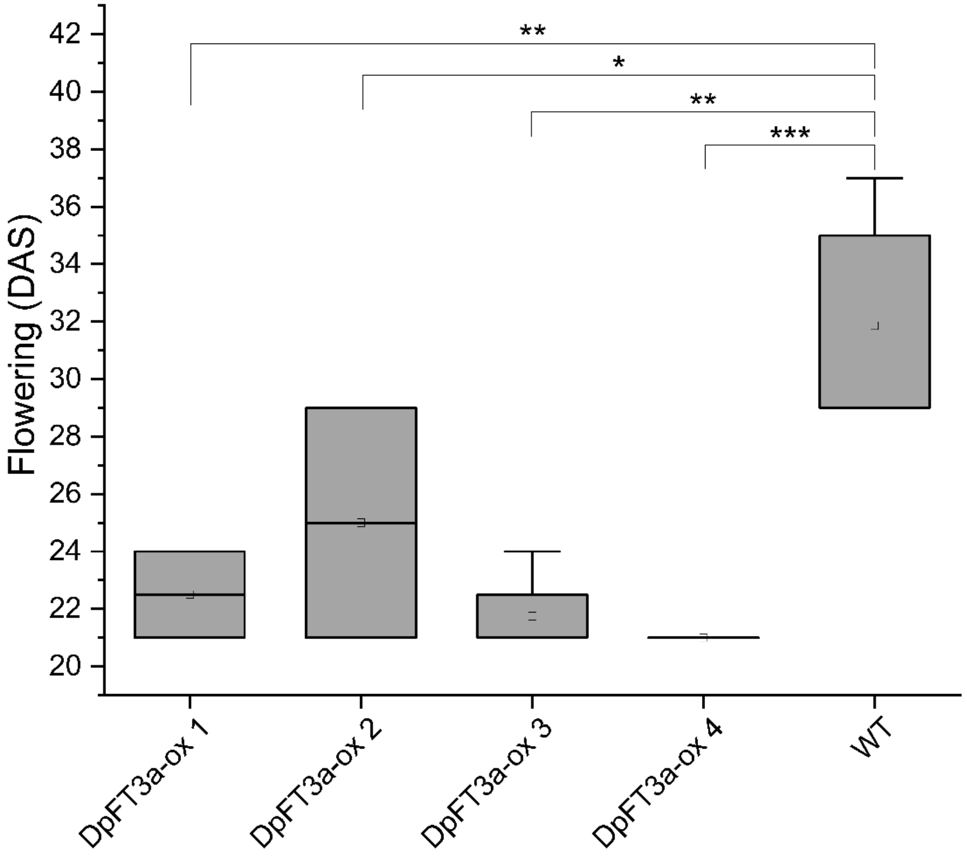


**Fig. S4.** Flowering time of DpFT3a overexpression lines. Overexpression of DpFT3a in four individual lines (DpFT3a-ox, n=4) of transgenic *A. thaliana* Col-0 plants compared to wild type (WT, n=6) plants. Flowering time in days after sowing (DAS). Difference in flowering time between lines was calculated via Wilcoxon ranked sum test (n=4). Asterisks indicate statistical significance: * p-value ≤ 0.05, ** p-value ≤ 0.01, *** p-value ≤ 0.001.

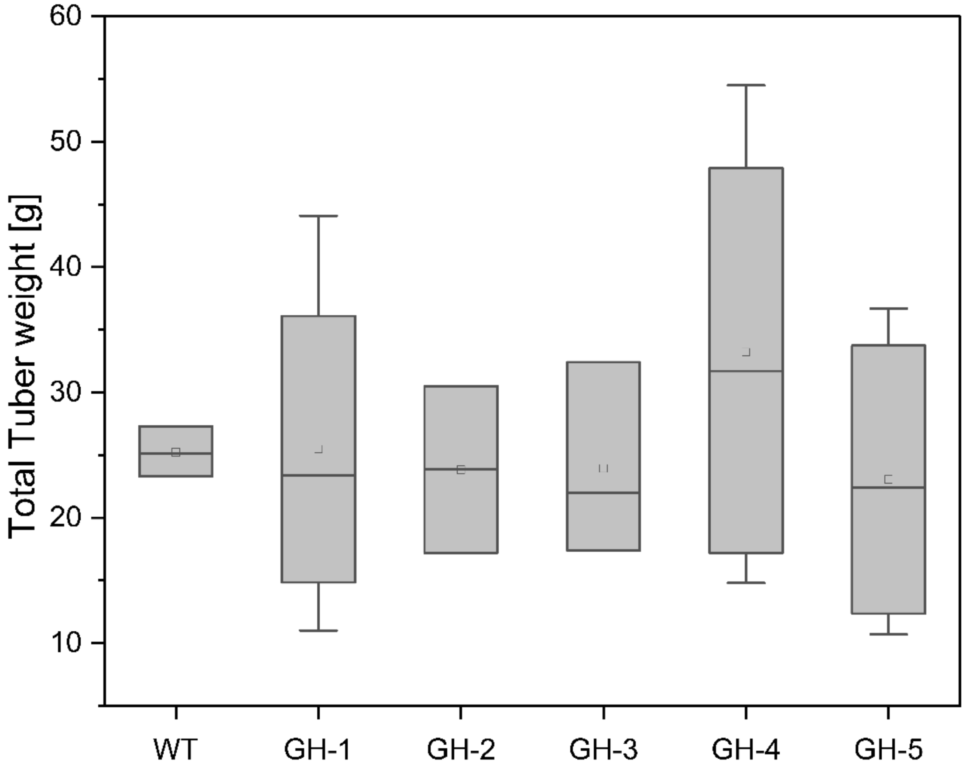


**Fig. S5.** Total tuber weight of ripe transgenic potato lines with *DpFT3a* overexpression and wildtype plants. Difference in the weight of harvested tubers between lines was calculated via Wilcoxon ranked sum test (n=2-5). Asterisks indicate statistical significance: * p-value ≤ 0.05, ** p-value ≤ 0.01, *** p-value ≤ 0.001

**Table S2.** Primers used in this study.

| Primer | Sequence (5'-3') | Purpose |
|---|---|---|
| DpFT3_NcoI_fwd | AAACCATGGGATGGAGGAACTTGATGGATAG | Amplification of DpFT3a for the generation of transgenic plants |
| DpFT3_XbaI_rev | AAACTCGAGTTACCTTCTACCGCCACAA | Amplification of DpFT3a for the generation of transgenic plants |
| DpFT3_qPCR1_fwd | AGTTAATGAGCCAAGGGTGGA | qPCR |
| DpFT3_qPCR1_rev | TCTGGTATATCTGTCACCAACCAAT | qPCR |
| qRT_StGAPDH_fw | ATGGCCTTCAGAGTACCAACTG | qPCR |
| qRT_StGAPDH_rev | ATGCTTGACCTGCTGTCACCAA | qPCR |
| Act42 | CCTCATCATACTCGGCCTTGGAG | semi-quantitative PCR |
| Act44 | GTAAGAGACATCAAGGAGAAGCTCTC | semi-quantitative PCR |
| DpFD1_NcoI-ATG_fwd | AAACCATGGGAGCATCGCCGCGGG | Amplification of DpFD1 for the generation of transgenic plants |
| DpFD1_XhoI-Stop_rev | AAACTCGAGCTACCATTGGGCGGAGC | Amplification of DpFD1 for the generation of transgenic plants |

**Table S2: Transcripts per Million**

| Gene ID | Tuber Head | Tuber Middle | Tuber Tip | DpPEBP Name |
|---|---|---|---|---|
| d_polystachya_ptg000024l_000055.1 | 0 | 0 | 0 | DpFT1a |
| d_polystachya_ptg000164l_000049.1 | 0 | 0 | 0 | DpFT1b |
| d_polystachya_ptg000166l_000157.1 | 0 | 0.023503192 | 0 | DpFT1c |
| d_polystachya_ptg000024l_000054.1 | 0 | 0 | 0 | DpFT2a |
| d_polystachya_ptg000164l_000048.1 | 0 | 0 | 0 | DpFT2b |
| d_polystachya_ptg000166l_000156.1 | 0 | 0 | 0 | DpFT2c |
| d_polystachya_ptg000008l_000149.1 | 5.56601681 | 2.107299892 | 1.968734905 | DpFT3a |
| d_polystachya_ptg000008l_000151.1 | 2.198356579 | 0.761409623 | 0.970698377 | DpFT3b |
| d_polystachya_ptg000058l_000117.1 | 13.4377811 | 5.796277477 | 3.741693014 | DpFT3c |
| d_polystachya_ptg000058l_000119.1 | 0 | 0 | 0 | DpFT3d |
| d_polystachya_ptg000079l_000508.1 | 5.911627355 | 1.943138775 | 1.376203633 | DpFT3e |
| d_polystachya_ptg000079l_000512.1 | 0 | 0 | 0 | DpFT3f |
| d_polystachya_ptg000086l_000448.1 | 6.844240105 | 2.734942854 | 3.462906604 | DpFT3g |
| d_polystachya_ptg000106l_000674.1 | 9.106847285 | 3.282125059 | 2.054176303 | DpFT3h |
| d_polystachya_ptg000106l_000676.1 | 0 | 0 | 0 | DpFT3i |
| d_polystachya_ptg000078l_000546.1 | 0 | 0 | 0 | DpFT4a |
| d_polystachya_ptg000044l_000186.1 | 0 | 0 | 0 | DpFT4b |
| d_polystachya_ptg000005l_000212.1 | 0 | 0 | 0 | DpFT4c |
| d_polystachya_ptg000031l_000159.1 | 2.831591388 | 0.065643081 | 0.121017864 | DpFT5a |
| d_polystachya_ptg000196l_000382.1 | 8.344418394 | 0.110272501 | 0.20957242 | DpFT5b |
| d_polystachya_ptg000104l_000205.1 | 0.854843489 | 0.007822825 | 0 | DpFT5c |
| d_polystachya_ptg000089l_000192.1 | 3.734119483 | 0.06842753 | 0.06935875 | DpFT5d |
| d_polystachya_ptg000012l_000528.1 | 0 | 0 | 0 | DpFT6a |
| d_polystachya_ptg000160l_000562.1 | 0 | 0 | 0 | DpFT6b |
| d_polystachya_ptg000100l_000524.1 | 0.029191019 | 0.015248361 | 0.016081309 | DpFT6c |
| d_polystachya_ptg000078l_000545.1 | 88.82247246 | 25.54145538 | 17.57456939 | DpFT7a |
| d_polystachya_ptg000044l_000185.1 | 73.49957201 | 18.83951067 | 11.1810326 | DpFT7b |
| d_polystachya_ptg000005l_000210.1 | 65.49918436 | 15.83179759 | 13.22921662 | DpFT7c |
| d_polystachya_ptg000063l_000722.1 | 0.40499937 | 2.212122081 | 0.767759195 | DpFT8a |
| d_polystachya_ptg000004l_000199.1 | 0.077467406 | 0.931032208 | 0.341898497 | DpFT8b |
| d_polystachya_ptg000093l_000706.1 | 0.301323146 | 0.861090861 | 0.323355299 | DpFT8c |
| d_polystachya_ptg000011l_000447.1 | 0.283613826 | 0.394215169 | 0.043181804 | DpFT9a |
| d_polystachya_ptg000010l_000250.1 | 0.030936641 | 0.125127467 | 0 | DpFT9b |
| d_polystachya_ptg000074l_000481.1 | 0.312262187 | 0.300304383 | 0.021152592 | DpFT9c |
| d_polystachya_ptg000024l_000239.1 | 0.290843903 | 0.190383491 | 0.011936673 | DpMFT1a |
| d_polystachya_ptg000164l_000241.1 | 0.76042309 | 0.320960899 | 0.008829242 | DpMFT1b |
| d_polystachya_ptg000180l_000084.1 | 0.296502474 | 0.077993125 | 0.024734085 | DpMFT1c |
| d_polystachya_ptg000003l_000697.1 | 0 | 0 | 0 | DpMFT2a |
| d_polystachya_ptg000057l_000739.1 | 0 | 0 | 0 | DpMFT2b |
| d_polystachya_ptg000066l_000779.1 | 0 | 0 | 0 | DpMFT2c |

| | | | | |
|---|---|---|---|---|
| d_polystachya_ptg000168l_000374.1 | 0 | 0 | 0 | DpMFT2d |
| d_polystachya_ptg000168l_000373.1 | 0 | 0 | 0 | DpMFT2e |
| d_polystachya_ptg000168l_000371.1 | 0 | 0 | 0 | DpMFT2f |
| d_polystachya_ptg000181l_000335.1 | 0 | 0 | 0 | DpMFT2g |
| d_polystachya_ptg000003l_000699.1 | 0 | 0 | 0 | DpMFT2h |
| d_polystachya_ptg000181l_000336.1 | 0 | 0 | 0 | DpMFT2i |
| d_polystachya_ptg000003l_000694.1 | 0 | 0 | 0 | DpMFT2j |
| d_polystachya_ptg000003l_000695.1 | 0 | 0 | 0 | DpMFT2k |
| d_polystachya_ptg000003l_000696.1 | 0 | 0 | 0 | DpMFT2l |
| d_polystachya_ptg000057l_000741.1 | 0 | 0 | 0 | DpMFT2m |
| d_polystachya_ptg000066l_000778.1 | 0 | 0 | 0 | DpMFT2n |
| d_polystachya_ptg000066l_000780.1 | 0 | 0 | 0 | DpMFT2o |
| d_polystachya_ptg000066l_000781.1 | 0 | 0 | 0 | DpMFT2p |
| d_polystachya_ptg000168l_000372.1 | 0 | 0 | 0 | DpMFT2q |
| d_polystachya_ptg000181l_000334.1 | 0 | 0 | 0 | DpMFT2r |
| d_polystachya_ptg000057l_000738.1 | 0 | 0 | 0 | DpMFT2s |
| d_polystachya_ptg000031l_000309.1 | 0 | 0 | 0 | DpTFL1a |
| d_polystachya_ptg000089l_000349.1 | 0 | 0 | 0 | DpTFL1b |
| d_polystachya_ptg000104l_000374.1 | 0 | 0 | 0 | DpTFL1c |
| d_polystachya_ptg000196l_000539.1 | 0 | 0 | 0 | DpTFL1d |
| d_polystachya_ptg002384l_000005.1 | 0 | 0 | 0 | DpTFL1e |
| d_polystachya_ptg000058l_000293.1 | 0.197491752 | 0.857717194 | 2.681045151 | DpTFL2a |
| d_polystachya_ptg000070l_000492.1 | 0.334561893 | 0.774677243 | 1.618479814 | DpTFL2b |
| d_polystachya_ptg000079l_000699.1 | 1.024705403 | 1.136557751 | 2.667609608 | DpTFL2c |
| d_polystachya_ptg000106l_000863.1 | 0.22082323 | 0.53733667 | 1.026422107 | DpTFL2d |
| d_polystachya_ptg000003l_000492.1 | 0.163001521 | 0 | 0.148303078 | DpTFL3a |
| d_polystachya_ptg000010l_000007.1 | 0.250582241 | 0.256032529 | 0.209833099 | DpTFL3b |
| d_polystachya_ptg000122l_000069.1 | 0.873484402 | 0.013419311 | 0.550568992 | DpTFL3c |
| d_polystachya_ptg000181l_000169.1 | 0.071066436 | 0.013608816 | 0.391121942 | DpTFL3d |
| d_polystachya_ptg000005l_000341.1 | 3.665322155 | 0.576826314 | 1.310275962 | DpTFL4a |
| d_polystachya_ptg000078l_000662.1 | 7.593410195 | 2.68874719 | 2.051454276 | DpTFL4b |
| d_polystachya_ptg000003l_000064.1 | 0.355729764 | 0.105812865 | 0.037710798 | DpTFL5a |
| d_polystachya_ptg000057l_000284.1 | 0.157242517 | 0.069545079 | 0 | DpTFL5b |
| d_polystachya_ptg000066l_000262.1 | 0.613450502 | 0.10839229 | 0.085361435 | DpTFL5c |
| d_polystachya_ptg000126l_000785.1 | 0.713624795 | 0.172255283 | 0.093893067 | DpTFL5d |

**Supplementary Methods:**

**Generation and analysis of transgenic Arabidopsis plants**

The coding sequence of *DpFT3a* was amplified from cDNA of F60 tuber tissue using primers DpFT3_NcoI_fwd and DpFT3_XbaI_rev. Amplified fragments were purified using agarose gel electrophoresis and the Gel and PCR clean-up-Kit (Macherey-Nagel, Düren, Germany) and ligated into plant expression vector plab12.1-Q35S (Post et al. 2012) for overexpression in *A. thaliana* Col-0 resulting in plab12.1-35S-DpFT3a. The plab12.1-35S-DpFT3a was transformed into *A. thaliana* Col-0 using a modified version of the floral-dip method (Clough and Bent 1998). Transgenic plants of the T1 generation were cultivated in a growth chamber at 24 °C and long-day conditions and checked daily for the emergence of inflorescence.

**Generation and analysis of transgenic *Nicotiana benthamiana* plants**

For localization and interaction studies in *N. benthamiana*, the coding sequence of *DpFT3a* and *DpFD1*were amplified from cDNA of F60 tuber tissue using primers DpFT3_NcoI_fwd and DpFT3_XbaI_rev and DoFD-like2_NcoI-ATG_fwd DoFD-like2_XhoI-Stop_rev, respectively. Amplified fragments were purified using agarose gel electrophoresis and the Gel and PCR clean-up-Kit (Macherey-Nagel, Düren, Germany) and ligated into the Gateway entry vector pENTR4 vector (Thermo Fisher Scientific, Waltham, USA). For localization analysis, *DpFT3a* and *DpFD1* were integrated into the destination vector pBAtTL-Venus-ccdB (Müller et al. 2010) using LR clonase (Thermo Fisher Scientific, Waltham, Germany) via Gateway cloning technology resulting in pBatTL-Venus-DpFT3a and pBatTL-Venus-DpFD1, respectively. For bimolecular fluorescence complementation (BiFC) analysis, *DpFT3a* and *DpFD1* were integrated into the pBatTL-N-mRFP-ccdB and pBatTL-C-mRFP-ccdB (Jach et al. 2006) via Gateway cloning technology generating pBatTL-N-mRFP-DpFT3a and pBatTL-C-mRFP-DpFD1, respectively.

Constructs were transformed into *Agrobacterium tumefaciens* strain GV3101 pMP90 by electroporation and infiltrated into *N. benthamiana* leaves as previously described (Müller et al. 2010). Plant cultivation and confocal laser scanning microscopy (CLSM) analysis was performed as described previously (Beinecke et al. 2018) using a Leica TCS SP5X microscope with excitation/emission wavelengths of 543/564–626 nm for mRFP and 514/525–600 nm for Venus.

**Literatur:**

Beinecke FA, Grundmann L, Wiedmann DR, Schmidt FJ, Caesar AS, Zimmermann M, Lahme M,

Twyman RM, Prüfer D, Noll GA (2018) The FT/FD-dependent initiation of flowering under long-day conditions in the day-neutral species Nicotiana tabacum originates from the facultative short-day ancestor Nicotiana tomentosiformis. Plant J 96:329–342. https://doi.org/https://doi.org/10.1111/tpj.14033

Clough SJ, Bent AF (1998) Floral dip: a simplified method for Agrobacterium-mediated transformation of Arabidopsis thaliana. Plant J 16:735–743. https://doi.org/10.1046/j.1365-313x.1998.00343.x

Jach, G., Pesch, M., Richter, K., Frings, S. and Uhrig, J.F. (2006) An improved mRFP1 adds red to bimolecular fluorescence complementation. Nat. Methods, 3, 597–600.

Müller B, Noll GA, Ernst AM, Rüping B, Groscurth S, Twyman RM, Kawchuk LM, Prüfer D (2010) Recombinant artificial forisomes provide ample quantities of smart biomaterials for use in technical devices. Appl Microbiol Biotechnol 88:689–698. https://doi.org/10.1007/s00253-010-2771-4

Post J, van Deenen N, Fricke J, Kowalski N, Wurbs D, Schaller H, Eisenreich W, Huber C, Twyman RM, Prüfer D, Gronover CS (2012) Laticifer-specific cis-prenyltransferase silencing affects the rubber, triterpene, and inulin content of Taraxacum brevicorniculatum. Plant Physiol 158:1406–1417. https://doi.org/10.1104/pp.111.187880

Müller, B. et al. Recombinant artificial forisomes provide ample quantities of smart biomaterials for use in technical devices. Appl. Microbiol. Biotechnol. 88, 689–698 (2010).
